# Secreted Clever-1 Modulates T Cell Responses and Impacts Cancer Immunotherapy Efficacy

**DOI:** 10.1101/2024.10.23.619796

**Authors:** Stuart Prince, Miro Viitala, Riikka Sjöroos, Ábris Á. Bendes, Jenna H. Rannikko, Daniel A. Patten, Ilaria di Benedetto, Laura Tyni, Carlos R. Figueiredo, Ilkka Koskivuo, Tiina A. Salminen, Shishir Shetty, Maija Hollmén

**Author notes:** Corresponding author: Maija Hollmén, PhD, MediCity Research Laboratory, University of Turku, Tykistökatu 6A, 20520 Turku, Finland.

## Abstract

Clever-1 functions as a scavenger and adhesion receptor, promoting tolerogenic activities in macrophages and subsets of endothelial cells, thereby contributing to cancer progression. High Clever-1 expression associates with resistance to immune checkpoint inhibitors and combined targeting of Clever-1 with anti-PD-1 enhances response in refractory mouse tumor models. A Clever-1–targeting humanized IgG4 antibody, bexmarilimab, is investigated in clinical trials as a macrophage-reprogramming therapy to treat solid tumors (NCT03733990) and hematological malignancies (NCT05428969). Here we describe a secreted form of (s)Clever-1 enriched in plasma of cancer patients, that was decreased upon bexmarilimab treatment. With the production of a recombinant sClever-1, mimicking the one found in human plasma, we show that sClever-1 can selectively bind activated T cells and disrupt T cell receptor signalling leading to impaired Th1 expansion. We demonstrate that sClever-1 binds to insulin growth factor 2 receptor (IGF2R) on T cells via its mannose-6-phosphate modification and further show that sClever-1 contributes to the immunosuppressive properties of macrophage-secreted extracellular vesicles, driving T cell tolerance and impairing anti-PD-1 efficacy. These findings suggest that Clever-1 exerts a systemic immunosuppressive effect independently of the cells it is expressed on, highlighting its potential as a target in cancer immunotherapy and a valuable biomarker for disease detection.

## Introduction

In recent years, the contribution of scavenger receptors to regulating macrophage responses has attracted significant attention. Scavenger receptors form a vital part of host defense and homeostasis by removing extraneous or modified self- and non-self-macromolecules and acting as co-receptors in the priming of effector immune responses when danger signals are detected. Common lymphatic endothelial and vascular endothelial receptor-1 (Clever-1, also known as Stabilin-1 and FEEL-1) is a multifunctional molecule that contributes to scavenging in a subset of some of the most anti-inflammatory macrophages. In these cells, Clever-1 is involved in receptor-mediated endocytosis and recycling, intracellular sorting, and transcytosis of altered and normal self-components (1). Clever-1 is upregulated when immunosuppression is needed to protect the body. For instance, during pregnancy, the immunosuppressive milieu that prevails in the placenta to support the growing fetus is associated with a high abundance of Clever-1-positive macrophages (2). Similarly, a growing tumor harnesses the immune system for its own development and is enriched with Clever-1-positive macrophages that support the formation of an immunosuppressive tissue microenvironment (3). Mechanistically, Clever-1 inhibits macrophage pro-inflammatory cytokine secretion and antigen presentation, thereby suppressing anti-tumor immunity mediated by CD8^+^ T cells (4,5). These observations have led to the clinical development of a humanized Clever-1-targeting antibody (bexmarilimab, FP-1305) to induce macrophage-mediated immune stimulation in various solid tumors (MATINS, NCT03733990) (6). A notable immunological finding from the MATINS trial is that anti-Clever-1 treatment induces robust peripheral T cell activation in patients with advanced cancer (5).

According to Adachi and Tsujimoto (7) there are two isoforms of Clever-1 (FEEL-1), one being the full-length (aa 2570) protein and the other being isoform #2, which has been predicted from cDNA and named soluble FEEL-1 due to lack of a transmembrane domain. The isoform #2 has not been verified to be expressed as protein but its cDNA sequence, ranging from the N-terminal part until the EGF-like domain 5 (aa 803), indicates that bexmarilimab (8) would also be able to bind this isoform. While it has been demonstrated that genetic silencing or antibody-mediated blockade of Clever-1 in human monocytes can promote IFNγ production in T cell antigen recall assays (4) it is still not clear whether Clever-1 itself has direct tolerogenic potential or these effects are mediated via an increase in pro-inflammatory monocyte activity during Clever-1 blockade. Our previous studies using placenta-purified Clever-1 protein show that it can directly bind T and B cells isolated from mouse spleen (9). However, the ligand for this interaction has not been determined.

While monocytes and macrophages are considered the main targets of bexmarilimab-induced anti-tumor responses (10), we describe here a secreted form of Clever-1 (sClever-1) in human plasma. In this study we explore the nature of this protein and show that it is enriched in plasma of cancer patients. By producing a recombinant form of a C-terminally truncated half Clever-1 molecule (H1) mimicking sClever-1 we show that it can bind activated T cells via cell surface IGF2R exposed at the same time point as transient phosphatidylserine (PS) exposure during early T cell activation (11) and drive their phenotype towards FoxP3 positive suppressive cells. Furthermore, our data also indicate that immunotherapeutic targeting of Clever-1 by bexmarilimab (Bex) in cancer patients can decrease the release of sClever-1 and reduce sClever-1 engagement with CD8^+^ T cells that was found to be important for the suppressive effect of macrophage-secreted extracellular vesicles.

## Materials and Methods

### Human samples

Blood samples were obtained from treatment naïve breast cancer patients prior and three weeks after undergoing mastectomy at the Turku University Hospital between years 2017-2022. Patients having a tumor nodule exceeding 2 cm in diameter were included in the study. Neoadjuvant therapy was not used. The EDTA-plasma was separated by centrifugation (2,000*g* for 10 min) within two-hours of withdrawal and stored at -70°C for later use. Peripheral blood mononuclear cells (PBMC) were isolated from the remaining EDTA-blood by Ficoll-Paque centrifugation and stored in freezing buffer (RPMI, 10% FCS, 1% glutamine, penicillin/streptomycin and 10% DMSO) at - 150°C for later use. The collection was done under the license ETMK: 132/2016 or 34/1801/2021 and approved by the ethical committee of the Turku University Hospital district. A written informed consent was obtained from each patient.

MATINS pre-treatment blood samples were collected per study protocol and centrifuged (as above) within 24 hours of withdrawal to obtain heparin plasma. Written informed consent was obtained from all study participants. The full study details can be found in Rannikko *et al.*(10).

All human liver tissue samples were obtained with prior written informed consent and were ethically approved for use in research. Normal liver tissues were obtained from rejected donor organs and chronically diseased liver tissues were collected from patients undergoing transplantation at the Queen Elizabeth Hospital, Birmingham, UK. All tissues were collected and used in accordance with the regulations and guidelines sanctioned by the West Midlands – South Birmingham Research Ethics Committee, Birmingham, UK (LREC reference no. 18/WA/0214).

Lymph was collected aseptically (needle aspiration) via postoperative lymphatic leakage at the Department of Otorhinolaryngology-Head and Neck Surgery, Turku University Hospital. The lymph was cleared by centrifugation (2,000*g* for 10 min) within two-hours of withdrawal and stored at -70°C for later use. The collection was done under the license T06/033/18 and approved by the ethical committee of the Turku University Hospital district. A written informed consent was obtained from each patient.

Blood and cells from healthy donors were collected under the license ETMK: 43/1801/2015 after receiving written informed consent. The collection was approved by the Turku University Hospital district.

Serum samples from lung carcinoma (n = 42) and melanoma (n = 16) patients treated with anti-PD-1 or anti-PD-L1 were obtained from Auria Biobank (Turku, Finland) with a dataset including information on dosing frequency, patient time on immunotherapy, blood cell counts and other treatments during or after immunotherapy (chemotherapy, radiotherapy and operations).

### Time-resolved fluorescence immunoassay (TRFIA)

A detailed description of the development of a sandwich-based ELISA assay using two different anti-Clever-1 antibodies with europium detection can be found in the supplemental material.

### Age as a contributing factor for plasma sClever-1

To evaluate the relationship between plasma sClever-1 levels and age in healthy controls, breast cancer patients and MATINS trial patients, sClever-1 levels were ln-transformed to make data conform to normal distribution. Pearson’s correlation coefficients and significances of correlation were calculated using R (v4.0.4, function cor.test). In addition, the relationship between ln-transformed sClever-1 and age was visualized separately in each MATINS trial cancer cohort using a scatter plot and a least-squares-fitted line. For predicting the effects of cancer and age on plasma sClever-1 levels, we performed multiple linear regression for a dataset consisting of treatment-naïve breast cancer patients and healthy controls. The following two models were fitted in the data (R function lm):

1. ln(sClever-1_i_) = β_0_ + β_1_ × Age_i_ + β_2_ × Cancer_i_ + ε_i_
2. ln(sClever-1_i_) = β_0_ + β_1_ × Age_i_ + β_2_ × Cancer_i_ + β_3_ × Age_i_ × Cancer_i_ + ε_i_

where 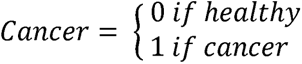

The two fitted models were compared with an F-test test (R function ANOVA, H_0_: β_3_= 0), which revealed that the model 2 was not improved by the addition of the interaction term (*P*-value = 0.8839), and therefore we report the model 1. By visual inspection of residual plots, we confirmed data linearity, residual homoskedasticity and roughly normal distribution of residuals.

### Primary cells

Peripheral blood mononuclear cells were separated by Ficoll-Paque PLUS (GE) density gradient centrifugation from which monocytes were enriched using CD14 magnetic beads (Human Monocyte Isolation Kit human, Miltenyi Biotec, RRID:AB_2665482) and T cells using Human T Cell Isolation Kit (negative selection, Stemcell). The monocytes were differentiated into macrophages by a seven-day incubation in IMDM (Gibco) supplemented with 10% FCS (Merck), penicillin/streptomycin (P/S) (Thermo Fisher) and 50 ng/mL of M-CSF (Biolegend) with one medium change. At this point the differentiated cells are referred to as M0. The macrophages were then polarized to M1 by first adding 20 ng/mL of recombinant IFNγ for 24h and then 100 ng/mL of LPS (Invivogen) for another 24h. For M2 polarization, the macrophages were incubated with recombinant human IL-4 (Peprotech) and 100 nM of dexamethasone (Merck) for 48h.

Human lymphatic endothelial cells (HLEC, No.2500) were purchased from ScienCell and cultured in Endothelial Cell Medium (ECM, No.1001). Human pulmonary microvascular endothelial cells (HPMEC, C-12281) were purchased from PromoCell and cultured in Endothelial Cell Growth Medium (ECGM, #C-22020) with supplement mix (#C-39225, both from PromoCell).

### LSEC isolation and stimulation

Liver sinusoidal endothelial cells (LSECs) were isolated from ∼75 g human liver tissue as described previously(12). Briefly, tissues were subjected to mechanical dissociation with scalpels and then enzymatic digestion with 10 mg/ml collagenase IA (Sigma-Aldrich). Resultant cell suspensions were separated out via density gradient centrifugation on a 33% / 77% Percoll (GE Healthcare) gradient at 850*g* for 25 min, brake 0. The non-parenchymal cell layer was then removed and LSECs were isolated via positive selection using CD31 antibody-conjugated Dynabeads^TM^ (Invitrogen, RRID:AB_3661738). LSECs were resuspended in human endothelial serum-free medium (SFM; Invitrogen), supplemented with 10% human serum (HD Supplies), 10 ng/mL vascular endothelial growth factor (VEGF; PeproTech), and 10 ng/mL hepatocyte growth factor (HGF; PeproTech), and seeded in a T-25 cell culture flask (Corning) pre-coated with rat tail collagen (RTC; 1 in 100; Sigma-Aldrich). LSECs were then cultured at 37°C in a humidified incubator with 5% CO_2_.

LSECs were seeded in RTC-coated 6-well tissue culture plates (Corning) and grown to confluence overnight. Cells were then stimulated with 2 mL complete LSEC medium containing 10 ng/mL tumor necrosis factor (TNF)-α and 10 ng/mL interferon (IFN)-γ for 24 hours. Supernatants were collected and centrifuged at 300*g*, before freezing at -20 °C until undergoing analysis for sClever-1.

### Cell lines

KG-1 (CCL-246, ATCC, RRID:CVCL_0374) were cultured in IMDM supplemented with 20% FCS and P/S. To induce differentiation into macrophages they were incubated in 100 ng/mL of phorbol-myristate acetate for 72h. The other cell lines used are listed with their specific culture medium: HEK293T (CRL-1573, ATCC, RRID:CVCL_0045; DMEM + 1% FCS); Jurkat (ATCC, RRID:CVCL_0367), Jurkat-Dual^TM^, Jurkat-Lucia^TM^ TCR-h-PD-1, Raji-APC-hPD-L1 (all from InvivoGen, RRID:CVCL_X593; IMDM, 2 mM L-glutamine, 25 mM HEPES, 10% FCS, P/S, 100 µg/mL Normocin).

### Separation of sClever-1 fractions by ultracentrifugation

KG-1 acute myelogenous leukemia cells were used as a positive control for high Clever-1 expression. Culture supernatant from 50-70% confluent KG-1 cells was collected. Blood and lymph were first diluted with an equal volume of PBS (1:1) due to their viscosity. All samples were then centrifuged by the optimized routine proposed by Théry *et al.*(13), without the filtration process for exosomes purification(13). Briefly, the samples were first centrifuged at 300*g* for 10 min at 4°C using Sorvall RT6000B refrigerated centrifuge for the removal of cells. Then cells were discarded, and the supernatant was collected and centrifuged at 2,000*g* for 10 min at 4°C using Sorvall RT6000B refrigerated centrifuge for the removal of cell debris. The supernatant was then collected and centrifuged at 20,000*g* for 30 min at 4°C using rotor SS-34, Sorvall RC5C centrifuge. The MVs pellet was then collected and suspended in 100-200 μL of PBS and stored at -20°C. For purifying the exosomes, the supernatant was transferred to ultracentrifuge tubes and then centrifuged at 100,000*g* for 70 min at 4°C using Beckman coulter, Optima L-90K ultracentrifuge. The exosome fraction was collected and the remaining supernatant was considered to contain free soluble Clever-1.

### Isolation of vesicle fractions

Plasma samples were pre-cleared by a two-step centrifugation, first at 2,000*g* for 20 min to remove cells and next at 10,000*g* for 20 min to remove debris. The plasma was diluted in PBS (1:1), mixed with Exosome Precipitation Reagent (from plasma, Invitrogen) and vortexed until the solution was homogenous. The samples were incubated at RT for 10 min and centrifuged at 10,000*g* for 5 min.

Cell conditioned media was pre-cleared by centrifugation at 2,000*g* for 30 min and a 0.5 volume of Exosome Precipitation Reagent (from cell culture media, Invitrogen) was added and vortexed to a homogenous solution. The samples were then incubated 4°C o/n and centrifuged at 10,000*g* for 1h at 4°C. Cold PBS was added and the samples were centrifuged at 14,000*g* for 5 min at 4°C. The supernatant was collected (liquid fraction) and the pellets were resuspended in PBS (vesicle fraction).

### Western blot

T cells were pelleted and resuspended in lysis buffer (10 mM Tris-HCl [pH 7.4], 5 mM NaF, 1.0% Triton X, 1 mM EDTA, 1 mM PMSF) supplemented with EDTA-free cOmplete Protease Inhibitor Cocktail (Roche). The cells were lysed at 4°C with mixing for 60 min and centrifuged at 14,000*g* and 4°C for 20 min. The supernatant was collected and Laemmli sample buffer (reducing) was added to the samples. The samples were run on a 4-20% FastGene PAGE gel (Nippon Genetics Europe). The gel was transferred to a PVDF membrane using the Trans Blot Turbo (Bio-Rad) protein transfer system. The membrane was blocked with 5% BSA in 0.1% Tween-TBS (Tris-buffered saline) and incubated sequentially with anti-pLck (cloneY394; R&D), anti-Lck (Cell Signaling) and anti-GAPDH (clone 6C5. HyTest) with in between stripping (1.5% Glycine, 0.1% SDS and 1% Tween 20, pH 2.2, 15 min at RT). The membrane was detected with ECL™ (Cytiva). IGF2R was detected using rabbit polyclonal anti-IGF2R antibody (20253-1-AP, Proteintech). For detection of mannose-6-phosphate and sClever-1, the samples were denatured at 70°C for 10 minutes but not reduced. Mannose-6-phosphate was detected by incubation with 10 μg/mL recombinant human IGF2R-His-AviTag protein (IGF-HM12RB, Kactus Bio) followed by 1 μg/mL HRP-conjugated rabbit anti-AviTag polyclonal antibody (orb475166, Biorbyt, RRID:AB_3661732). To detect sClever-1 the membrane was incubated with 1 μg/mL rat IgG2a monoclonal anti-human Clever-1 antibody 9-11 (InVivo Biotech, RRID:AB_3661734) or mouse anti-human Clever-1 antibody 4G9 (Santa Cruz Biotechnology, RRID:AB_3661731). When necessary, HRP was directly conjugated to the primary detection antibody, as in the case of anti-Clever-1-HRP (9–11), by using a HRP Conjugation Kit (ab102890, AbCam). For primary macrophage and EV blots, TFRC NBP2-34602 (Novus, RRID:AB_3289338), CD206 12981S (Cell Signaling), CD63 sc-5275 (Santa Cruz, RRID:AB_627877) were used.

### Pulldown of sClever-1 from serum

Human IgG4, bexmarilimab (both from Abzena, RRID:AB_3661735, RRID:AB_3661733), rat IgG2a isotype control (BioXcell, RRID:AB_1107769) and anti-Clever-1 9-11 (InVivo Biotech) antibodies were diluted to 1 mg/mL in PBS (100 µL) and biotinylated with NHS-PEG4-biotin (EZ-Link™, No-Weigh™ Format, Thermo) at a calculated ratio of 6 biotin molecules per molecule of antibody, according to the manufacturer’s instructions. Following the biotinylation reaction the excess biotin was removed by overnight dialysis in PBS in Slide-a-Lyzer MINI Dialysis Devices 10 kDa MWCO (Thermo).

Human AB serum (Millipore) was diluted 1/10 to a final volume of 200 mL in ice-cold PBS + 0.1% Tween 20. This was precleared by end-over-end mixing at 4°C overnight with 10 mg pre-washed Dynabeads M-280 Streptavidin (Invitrogen), divided equally into four 50 mL Falcon tubes. The beads were pelleted at 3000*g* and 4°C for 10 min, held in place with a magnet, and the precleared supernatant removed and divided into four separate 50 mL Falcon tubes on ice. Fifty micrograms of each biotinylated antibody were added separately to each tube of precleared supernatant. The biotinylated antibodies were mixed with the precleared supernatants by end-over-end mixing at 4°C for 8 h. Following this 2.5 mg of pre-washed Dynabeads M-280 Streptavidin were added to each tube to capture the complexes of biotinylated antibodies and ligands, and incubation was continued with end-over-end mixing at 4°C overnight. The following day the beads were pelleted at 3000*g* and 4°C for 10 min and the supernatant (“flow-through”) removed and frozen. The pelleted magnetic beads were resuspended in 1.5 mL of ice-cold PBS + 0.1% Tween 20, mixed thoroughly by vortexing then pulled-down using a DynaMag-2 magnetic rack (Invitrogen) and thoroughly aspirated. This washing procedure was repeated for a total of four times to remove non-specifically bound proteins from the beads. Finally, the fully aspirated beads were eluted in 40 µL SDS-PAGE denaturing sample buffer without reducing agent, pulled-down with the magnet, aspirated, and then the elution repeated in a second 40 µL volume of SDS-PAGE denaturing sample buffer without reducing agent. The 80 µL of eluate from each sample was pooled and heated at 70°C for 10 min prior to loading on an SDS-PAGE gel for Coomassie staining and isolation of specific bands for mass-spec proteomics analysis. For this purpose, 50 µL of sample was loaded per well of a 4-20% Mini-PROTEAN TGX precast gel (BioRad) and run to completion at 100 V. Spectra Multicolor High Range Protein Ladder (Thermo) was included as molecular weight standard. Following electrophoresis, the gel was washed in ddH20 and then stained in GelCode Blue Safe Protein Stain (Thermo) according to the manufacturer’s instructions. Following de-staining in ddH2O the gel was photographed before and after the excision of specific molecular weight bands of interest using a clean scalpel and collection tube. The collected bands were sent for mass-spec proteomics analysis. A repeat of the experiment for western blotting was also performed to confirm the presence and approximate molecular weight of Clever-1 pulled down. In a variation of this method, the magnetic Dynabeads used for the pulldown were washed 4 times in PBS with 0.1% Tween 20, then three times in PBS (to remove residual detergent) and sent for mass spectrometry and proteomics analysis of bound protein without any prior separation via SDS-PAGE. When necessary to detect mannose-6-phosphate on protein that was pulled down, the PBS was supplemented with 5 mM NaF, 1 mM EDTA, 1 mM PMSF and EDTA-free cOmplete Protease Inhibitor Cocktail (Roche).

### Expression, purification and modification of half Clever-1 (H1)

The codon-optimised cDNA of human Clever-1 (UniProt ID: Q9NY15) between residues 26-1255 was cloned into a pQMCF-1.2 expression vector between an N-terminal insulin signal peptide and a C-terminal His_8_-tag to create a 133.6 kDa soluble construct; H1. The recombinant expression and purification of H1 were performed by Icosagen Cell Factory. In summary, H1 was transiently expressed in CHOEBNALT85 cells using the QMCF technology(14). The secreted protein was captured on Ni^2+^ Sepharose affinity resin (Cytiva) and washed with high-salt buffer before elution with 0.3 M imidazole. A final polishing step was performed by size-exclusion chromatography (SEC) on a Superdex 200 Increase 10/300 GL (GE Healthcare) column using a 20 mM HEPES pH 8.0, 150 mM buffer NaCl, 2 mM CaCl_2_, and 5% glycerol buffer. This buffer (subsequently called modified HBS) is used for further experiments unless stated otherwise. The purity was assessed using Coomassie-stained SDS-PAGE, while end-point stability was tested before and after three rounds of freeze-thaw cycles by comparative analytical SEC on a BioSuite 250 4.6/300 column (Waters). Aliquots of H1 were snap-frozen for long-term -70°C storage and sterile-filtered using a 0.2 µm membrane before subsequent use.

To biotinylate H1 protein, “No Weigh” NHS-PEG4-Biotin (Thermo) was used to label recombinant H1 at a molar ratio of 6:1. The biotinylated protein was then dialyzed at 4°C in HBS to remove excess biotin. The dialyzed, biotinylated protein was filter-sterilized and aliquoted at -70°C. To measure how many molecules of biotin were conjugated per molecule of H1, a Pierce Biotin Quantitation Kit (Thermo) was used according to the manufacturer’s instructions. There were 3.8755 biotin molecules on average per molecule of H1. To determine what percentage of H1 molecules were biotinylated, a non-reducing denaturing SDS-PAGE gel was run with unheated non-reduced samples of H1, streptavidin, and H1 precomplexed with streptavidin, alongside Spectra Multicolor High Range Protein Ladder (Thermo). The gel was stained with GelCode Blue (Thermo), and the molecular weight-shift of complexed H1 analysed. More than 95% of H1 was biotinylated.

To dephosphorylate H1, the protein was digested with Fast AP Alkaline Phosphatase enzyme (Thermo) by diluting the protein to a final concentration of 0.1 mg/mL in a final volume of 100 μL of 1x reaction buffer with 10 units of Fast AP enzyme, followed by incubation at 37°C overnight. To detect dephosphorylation, the loss of binding to recombinant IGF2R protein (Kactus Bio) on Octet was confirmed. To confirm integrity of the protein, the concentration was checked by NanoDrop and a sample run on SDS-PAGE and stained with GelCode Blue (Thermo).

### Light scattering

The absolute molecular mass of H1 was determined using size-exclusion chromatography-coupled multi-angle light scattering (SEC-MALS). 150 µg of H1 was injected on a Superdex 200 Increase 10/300 GL (GE Healthcare) column and eluted at RT using 0.75 mL/min flow rate in modified HBS without glycerol. The light scattering signal was recorded using a miniDAWN TREOS II (Wyatt Technology, RRID:SCR_020895) detector. Online concentration was measured with an Optilab T-rEX (Wyatt Technology, RRID:SCR_020891) differential refractometer. Bovine serum albumin (Sigma-Aldrich) was used for calibration. The glycosylation level was derived using conjugate analysis in the ASTRA software (Wyatt Technology, RRID:SCR_016255) assuming refractive index increment values of 0.185 and 0.156 mL/g for the protein and the glycan component, respectively(15).

Hydrodynamic radius of the monomeric H1 was assessed using dynamic light scattering (DLS) on a Zetasizer Nano ZS (Malvern, RRID:SCR_025760) system. Data acquisition was carried out at RT with a protein concentration of 0.2 mg/mL. Six measurements were averaged to determine the intensity size distribution.

### Circular dichroism spectroscopy (CD)

CD spectra between 180-250 nm were recorded on a Chirascan CD spectrophotometer (Applied Photophysics, RRID:SCR_025759) at RT using a 0.1 mm path-length quartz cuvette. The measurement of H1 at a 1 mg/mL concentration was carried out in a buffer containing 20 mM sodium phosphate pH 8.0 and 150 mM NaF. Six scans were averaged and smoothed. As a reference, five representative models of H1 were prepared using AlphaFold2(16) (RRID:SCR_025454). Simulated CD spectra of the models were prepared and averaged using the PDBMD2CD server(17) (RRID:SCR_025768). The CD spectra were deconvolved(18) and classified(19) using the BeStSel server (RRID:SCR_025767)

### Differential scanning fluorimetry (DSF)

The stability of H1 was characterised by following its thermal unfolding at 0.2 mg/mL using a Prometheus NT.48 (NanoTemper, RRID:SCR_025758) system between 20-95°C with a ramping speed of 1 °C/min. The measurement was carried out in triplicates.

### Small-angle X-ray scattering (SAXS)

SEC-SAXS measurements were carried out on the CoSAXS beamline (RRID:SCR_025757) at the MAX IV synchrotron. 250 µg of H1 was applied on a Superdex 200 Increase 10/300 GL (GE Healthcare) column and resolved at RT with a 0.6 mL/min flow rate. SAXS data were recorded between scattering vectors 0.003-0.334 Å^-1^. The significant components in the elution data were reduced using regularized alternating least squares (REGALS) method(20) and analysed with BioXTAS RAW(21) (RRID:SCR_025769). The MW was derived from the whole q range using the Porod invariant method(22).

### T cell activation

T cell density was adjusted to 1.0 × 10^6^ cells/mL in RPMI 1640 medium (Sigma) supplemented with 10% FCS,100 U/mL penicillin, 100 µg/mL streptomycin, 55 µM 2-ME and 2 mM L-glutamine. The T cells were plated in 96-well plates (Sarstedt) at 1.0 × 10^5^ cells per well and activated either by ionomycin stimulation at 1 µg/mL (Sigma) or plate-coated anti-CD3 at 10 µg/mL (clone OKT3; Invitrogen, RRID:AB_468854) and anti-CD28 at 10 µg/mL (clone 28.2; Invitrogen, RRID:AB_468926) with IL-2 (Miltenyi Biotech) at 20 ng/mL. H1 or BSA control was added upon activation at 50 µg/mL for the whole duration of the experiment.

### Flow cytometry

T cells were activated with CD3/CD28+IL-2 for 48 h or ionomycin (up to 120 minutes) and thereafter incubated with 50 µg/mL of biotin-H1 (Icosagen) or biotin-IgG4 (Abzena) at 37°C for 30 min in PBS supplemented with 0.1% BSA followed by streptavidin-PE (5 µg/mL) (BD). When co-staining for Annexin V-FITC (10 µg/mL, BD) and FAM-VAD-FMK (1:250, AAT Bioquest) was performed, a Annexin V Binding Buffer containing 10 mM HEPES pH 7.4, 140 mM NaCl, 2.5 mM CaCl_2_ and 0.1% BSA was used. Recombinant Annexin V (10 µg/mL, BD) or H1 (50 µg/mL) at 37°C for 30 min was added before staining to test competitive binding to PS.

For measuring T cell proliferation, the cells were pre-labelled with 1 µM Vybrant™ CFDA SE dye (CFSE; Invitrogen) according to the manufacturer’s protocol and stained with anti-CD8-APC (clone RPA-T8, BD, RRID:AB_398595) after six days. For intracellular staining of lineage determining transcription factors, the cells were first surface stained with anti-CD8-BV510 (clone SK1, BD, RRID:AB_2722546), fixed and permeabilized using FoxP3/Transcription Factor Staining Buffer Kit (Thermo Fisher) according to the manufacturer’s protocol and stained with anti-FoxP3-PE (clone PCH101, Invitrogen, RRID:AB_1518782) and anti-T-bet-BV412 (clone 04-46, BD, RRID:AB_10564071). For assessing macrophage polarization after autologous T cell coculture cell surface staining was performed with the following antibodies: anti-CD64-PE (clone 10.1, BD, RRID:AB_1727085), anti-HLA-DR-PE-Cy7 (cloneG46-6, BD, RRID:AB_1727528), anti-CD163-BV711 (clone GH1/61, BD, RRID:AB_2738469), anti-CD40-BV510 (clone 5C3, BD, RRID:AB_2738218), anti-Clever-1-Alexa Fluor 647 (clone 9-11, in-house conjugated) or isotype control antibodies with the same fluorochromes.

For surface staining of Jurkat cells, the cells were pelleted and resuspended in ice-cold Annexin V Binding Buffer plus 100 μg/mL Kiovig (RRID:AB_3661740) and stained with biotinylated H1 as described for primary T cells. Following this, Annexin V-FITC was added for a 5-minute incubation at room temperature, where applicable. Thereafter, all washing, and incubation steps were performed on ice, in ice-cold Epics 1 Buffer (PBS, 2% FCS, 0.1% azide). Alexa Fluor 647 labelled streptavidin (Life Technologies) was used at 2 μg/mL, to detect H1 surface staining. Cells were finally fixed in 0.5% paraformaldehyde PBS before flow cytometry. For intracellular staining of Jurkat cells, the cells were first fixed in 4% paraformaldehyde in PBS, then permeabilized with PBS + 0.2% Tween 20. Thereafter, 0.1% Tween 20 was included in all steps to maintain permeability. Anti-TFRC (NBP2-34602; Novus) and anti-IGF2R (20253-1-AP; Proteintech, RRID:AB_10859779) were used to detect knockdown efficiency. The cells were analysed with LSRFortessa (BD, RRID:SCR_018655) and data processed with FlowJo software v.10.7.1 (TreeStar, RRID:SCR_008520). Pre-gating for viable cells and singlets was performed prior specific analyses.

Frozen aliquots of PBMCs from healthy donors and breast cancer patients were pre-stained with fixable viability dye eFluor 450 (Thermo) and thereafter with anti-IGF2R + goat anti-rabbit AF488 (Invitrogen, RRID:AB_143165), biotinylated H1 + PE-streptavidin (#349023, BD), anti-CD8-APC (clone RPA-T8, BD), anti-CD45RO-APC-H7 (clone UCHL1, BD, RRID:AB_10562194) and anti-PD-1-PerCP-Cy5.5 (EH12.1, BD, RRID:AB_10645786).

### Jurkat-Dual assay

Jurkat-Dual cells were plated 0.4 × 10^6^ cells/well in 96-well plates in 200 µL of cell culture medium supplemented with anti-CD3 and anti-CD28 as described above. H1 or BSA control was added upon activation at 50 µg/mL for 18 hours. NFκB activation was analysed by measuring the luciferase reporter from the cell culture supernatant with QUANTI-Luc reagent (InvivoGen) according to the manufacturer’s instruction using Tecan Infinite (Tecan).

### Cytokine multiplex

Primary T cell culture supernatants were analyzed for cytokine content six days after activation with Bio-Plex Pro™ Human Cytokine 27-Plex Assay (#M500KCAF0Y, BioRad) according to the manufacturer’s protocol.

### Retrogenix Cell Microarray assay

To discover H1 binding ligands, biotin-H1 was tested in the Retrogenix Cell Microarray assay consisting of 6449 human plasma membrane, tethered human secreted proteins (using an epitope tag and tethered single-chain variable fragment [scFv] anchor technology) and 397 heterodimer plasma membrane protein complexes (Charles River Discovery). For library screening 10 μg/mL of biotin-H1 was screened in duplicate for binding against fixed HEK293 cells using both the sequential method (test protein added to slides, washed and then AlexaFluor647 Streptavidin detection reagent added) and the pre-incubation method (test protein pre-incubated with AlexaFluor647 Streptavidin at a 4:1 molar ratio before addition to slides). Specific hits were confirmed by staining un-fixed and fixed cells and further validated by flow cytometry.

### Pulldown with biotinylated H1 in Jurkat cells and primary T cells

Method 1 (with crosslinker): Six million Jurkat cells were resuspended in 0.6 mL ice-cold Annexin V Binding Buffer with or without 50 μg/mL biotinylated H1 protein. The cells were then placed in a 37^°^C cell culture incubator for 30 minutes for H1 to bind to the cell outer membranes. The cells were then pelleted at 300*g* and washed twice with ice-cold PBS to remove unbound H1. The aspirated cell pellets were resuspended in freshly prepared 3.5 mM BS3 crosslinker (Pierce, No-Weigh Format) in 4^°^C PBS, then incubated for 30 minutes at room temperature for crosslinking. Twenty millimolar Tris pH 7.6 was added for 15 minutes to quench the crosslinker. The cells were then pelleted at 400*g*, aspirated, and washed twice with ice-cold PBS to remove residual unreacted crosslinker. The washed and pelleted cells were placed on ice and lysed in 1 mL ice-cold Lysis Buffer containing protease inhibitors. Following 15 minutes of lysis the cells were centrifuged at 4^°^C and 10,000*g* for 5 minutes to remove nuclei. The clarified supernatants were incubated with magnetic M280 Streptavidin Dynabeads (Invitrogen) overnight at 4^°^C with mixing, for pulldown of biotinylated H1 complexes to occur. Following the pulldown, the Dynabeads were pelleted and washed 4x in 1 mL ice-cold Lysis Buffer, then pelleted and washed 3x in TBS to remove residual detergent. The aspirated beads were then sent for mass spec and proteomics analysis to identify and quantify the proteins that had been co-precipitated with biotinylated H1.

Method 2 (with purified plasma membranes): Fifty million Jurkat cells were resuspended in 1.2 mL ice-cold Annexin V Binding Buffer with or without 50 μg/mL biotinylated H1 protein. After H1 binding the aspirated cell pellets were resuspended in 0.5 mL ice-cold PBS premixed 50:50 with hypotonic Buffer A from a Minute Plasma Membrane Protein Isolation Kit (Invent Biotechnologies) to which 1 mM PMSF, 5 mM NaF, Roche cOmplete EDTA-free Protease Inhibitor Cocktail had been added. The cell suspensions were incubated on ice for 5 minutes to sensitize the cells to rupturing by shear forces. The cell suspensions were then passed once through a Minute Plasma Membrane Protein Isolation Kit Filter Cartridge at 16,000*g* for 30 seconds to rupture the cells. Thereafter the isolation of plasma cell membranes and other separate cell fractions (nuclei, cytosol and organelles) were performed according to the manufacturer’s instructions. The final yield of plasma membrane protein was measured by micro–DC Protein Assay (Bio-Rad) and found to be between 55-57 μg for each sample. The purified plasma membrane pellets were solubilized in 0.5 mL ice-cold Lysis Buffer and streptavidin Dynabead pulldowns were performed as previously.

### Mass-spec, proteomics analysis and data filtering

Samples were on-bead digested according to standard protocol at the Turku Proteomics facility. Protein samples, bound on magnetic streptavidin beads, were denatured with 8 M urea in 50 mM Tris-HCl, pH 8. Proteins were reduced with 10 mM D,L-dithiotreitol (in 50 mM Tris-HCl, pH 8), and alkylated with 40 mM iodoacetamide (in 50 mM Tris-HCl, pH 8). Samples were digested overnight with sequencing grade modified trypsin (Promega). Peptide samples were desalted using Sep-Pak tC18 well plate (Waters) and peptides were dried in a vacuum centrifuge. Digested peptides were dissolved in 11 μL of 0.1% formic acid. From each digested peptide sample 5 μL was injected for analysis.

The LC-ESI-MS/MS analysis was performed on a nanoflow HPLC system (Easy-nLC1000, Thermo Fisher Scientific, RRID:SCR_014993) coupled to the Q Exactive HF mass spectrometer (Thermo Fisher Scientific, RRID:SCR_020425) equipped with a nano-electrospray ionization source. Peptides were first loaded on a trapping column and subsequently separated inline on a 15 cm C18 column (75 μm x 15 cm, ReproSil-Pur 3 μm 120 Å C18-AQ, Dr. Maisch HPLC GmbH, Ammerbuch-Entringen, Germany). The mobile phase consisted of water with 0.1% formic acid (solvent A) or acetonitrile/water (80:20 [v/v]) with 0.1% formic acid (solvent B). A 40 min long step gradient (from 5% to 21% of solvent B in 17 min, from 21% to 36% of solvent B in 13 min, and from 36% to 100% of solvent B in 5 min, followed by a 5 min wash stage with 100% of solvent B) was used to eluate peptides.

MS data was acquired automatically by using Thermo Xcalibur 4.1 software (Thermo Fisher Scientific, RRID:SCR_014593). An information dependent acquisition method consisted of an Orbitrap MS survey scan of mass range 350–1750 m/z with resolution of 120,000, a target value of 3,000,000, and maximal injection time of 100 ms. The 15 most intense peptide ions were selected for MS2 fragmentation with resolution of 15,000, a target value of 50,000, and injection time of 200 ms. Data files were searched for protein identification using Proteome Discoverer 3.0 software (Thermo Fisher Scientific, RRID:SCR_014477) connected to an in-house server running the Mascot 2.8.2 software (Matrix Science, RRID:SCR_014322). Variable modifications included where applicable were mass shift of hydrolyzed BS3 and the mass shift of Tris-quenched BS3. Data was searched against a SwissProt database (version 2022_03, RRID:SCR_013448) with taxonomy filter *Homo sapiens*.

For each experiment + control “pair”, the results of the experiment pulldown with biotinylated H1 were compared to the results of the control pulldown without any biotinylated H1. Any hits where the Score Mascot for the experiment was less than threefold greater than the Score Mascot for the control were filtered out to remove noise, such as for e.g., keratin, albumin, etc. Unused filters added for qualitative assessment were Retrogenix array score and CRAPome (https://reprint-apms.org, <u>RRID:SCR_025008</u>) score. The hits were ranked according to number of peptides for a given protein. Any hit protein with less than two unique peptides was deprioritized. The results of the pulldown with crosslinking were compared to the results for the pulldown with non-crosslinked purified plasma membranes. STRING database (https://string-db.org, <u>RRID:SCR_005223</u>) node analysis was performed to look for patterns in the results.

### Octet

Measurement of protein interactions via biolayer interferometry (BLI) was performed using a Fortebio Octet RED384 (RRID:SCR_023267) in 8-channel mode and 96-well plate format. All assays were performed at 1000 rpm, 25°C, and in 200 μL Kinetics Buffer (10 mM HEPES pH 7.4, 150 mM NaCl supplemented with 0.1% BSA and 0.02% Tween 20). The assay definition included baseline (5 min), loading (5 min), association (3 min), dissociation (30 min). The biosensors used were streptavidin (Sartorius). The biotinylated ligands loaded onto the streptavidin biosensors at 10 μg/mL were either biotinylated H1 protein or biotinylated IGF2R domains 1-10 protein (Kactus Bio), as applicable. The non-biotinylated solution analyte proteins used in binding assays were either H1 protein or IGF2R domains 11-13 (R&D Systems), as applicable. All experiments included a negative control reference biosensor for subtraction. Single cycle kinetics data was fitted using TraceDrawer software (RRID:SCR_025782) according to a 1:1 Langmuir model.

### siRNA and CRISPR/Cas9

Jurkat cells were electroporated with pooled Edit-R Predesigned sgRNAs targeting IGF2R (SG-010601-01, SG-010601-02, SG-010601-03) or TFRC (SG-003941-01, SG-003941-02, SG-003941-03; all from Dharmacon) complexed with TrueCut Cas9 Protein v2 using the SE Cell Line kit and 4D-Nucleofector X Unit (Lonza, RRID:SCR_023155) according to manufacturer’s protocol.

Primary human monocytes were positively enriched from buffy coat PBMCs with CD14 Microbeads and LS columns (Miltenyi Biotec). TrueCut Cas9 Protein v2 and TrueGuide Synthetic single guide (sg)RNAs (both from Thermo Fisher) were complexed at RT and delivered into purified monocytes by electroporation as described.(23) Subsequently, the monocytes were differentiated into macrophages as described above. The sgRNAs targeted the sequences ACGGAAGTGCCGGAAGCAAG (sg408) and GGGTACATGAGTGACAAACG (sg418) in the human *STAB1* gene. sgRosa26 (A35525) targeting the murine Rosa26 locus was used as a negative control.

KG-1 cells were diluted (2.0 × 10^6^ cells per electroporation) in OPTIMEM (Gibco™ Opti-MEM™ I Reduced Serum Medium 31985062). siRNAs were added to 2 µM and electroporation was performed using the program U-001 on the 3D-Nucleofector (Lonza). The siRNAs targeting Clever-1, AUGAUGAGCUCACGUAUAA (siR1) or UCAAGUCGCUGCCUGCAUA (siR4) (J-014103-05-0020 and J-014103-08-0020, respectively) as well as the negative control “scrambled” siRNA (ON-TARGETplus Control Pool D-001810-10-20) were ordered from Dharmacon. After electroporation the KG-1 cells were differentiated into macrophages by a three-day incubation with 100 nM phorbol-myristate-acetate (PMA). Cell supernatants were collected, cleared and EVs collected using the precipitation method. The collected EVs (2.5 μg/mL) were used in T cell activation assay as previously described.

### EV enrichment from conditioned media using size exclusion chromatography

For EV enrichment from conditioned media, floating cells were pelleted at RT and 300*g* for 10 min and larger vesicles at RT and 2,000*g* for 30 min. The conditioned medium was then concentrated from 15 mL to ∼200 µL using 100 kDa MWCO Amicon Ultra centrifugal filters (Millipore) and centrifuged at 4°C and 20,000*g* for 30 min. The supernatant was carefully removed and sterile filtered (200 nm). 150 µL of clarified supernatant was fractionated on the qEV Automatic Fraction Collector using 70 nm qEV Single columns (Izon, RRID:SCR_025764). The presence of EVs was determined flow cytometrically using Exosome-Human CD63 Isolation/Detection Reagent according to manufacturer’s instructions (Invitrogen) and the absence of contaminating soluble proteins using the Qubit Protein Assay (Invitrogen, RRID:SCR_020311). Fractions containing EVs were pooled, concentrated further and used for functional experiments.

### Jurkat-Lucia^TM^ TCR-h-PD-1 – Raji-APC-hPD-L1 assay

Jurkat-Lucia™ TCR-hPD-1 cells (0.2 × 10^6^) and Raji-APC-Null cells (0.1 × 10^6^) were mixed in round bottom clear wells (Greiner CELLSTAR® 96 well plates, M9311) and incubated for 30 min at 37°C. Thereafter 1 μg/mL Anti-hPD-1-Ni-hIgG4 and 0.5 μg of isolated EVs were added. The plate was incubated at 37°C in a CO_2_ incubator for 6 h. Luminescence signal was measured using QUANTI-Luc™ reagent and Tecan Infinity (TECAN, RRID:SCR_024560) with the following parameters: end-point measurement with a 4 sec start time and 0.1 sec reading time.

### Time/dose-dependent analysis of Clever-1 levels on circulating T cells and monocytes/macrophages (MoMacs)

To analyze PBMCs from patients stained with a CyTOF macrophage panel 2 (5), we implemented Hierarchical Stochastic Neighbor Embedding (HSNE) due to its multi-scale data handling, scalability to large datasets, and robustness to noise—critical for detecting subtle surface differences in rare populations (24), making it ideal for evaluating small variations in cell protein binding to immune cells post bexmarilimab treatment. Cytosplore version 2.3.1 (RRID:SCR_018330) was used.

The following phenotyping markers were used for clustering: CD16, CD4, KI67, CD11C, 9-11, CD3, CD8, CD19, CD56, and HLA-DR, with a hyperbolic transformation applied for standardization. The analysis focused on T-cell clusters (CD3^+^CD4^+^ and CD3^+^CD8^+^) and macrophage-like clusters identified as CD19^-^, CD3^-^, CD56^-^, CD16^-^, CD163^+^, and HLA-DR^+^. HSNE parameters in Cytosplore included the Euclidean transformation for KN metric, RNG seed of -1, memory-preserving computation, checks in AKNN set at 1,024, #RW for influence at 100, #RW Monte Carlo at 15, random walks threshold at 1.5, random walks length at 15, pruning thresholds at 0, filter percentile at 0.5, four scales, and cache HSNE hierarchy enabled without GPU acceleration. The analysis was performed on concatenated patient samples from each time point, ensuring normalization across all patients for each day. Mean intensity values of Clever-1 for T cell and macrophage clusters were computed, with heat map visualizations depicting expression levels from red (highest) to dark blue (lowest), facilitating clear comparative analysis. Each patient’s data contributed to a single mean value per cluster per time point. Data from all patients were consolidated, resulting in a readout for each day, with seven patients for day 1 and day 8, and six patients for day 15. Mean intensity values were visualized using bar plots, illustrating expression levels across different clusters and time points.

### Statistics

Data are presented as mean ±SD unless otherwise noted. Frequency distribution was calculated for sClever-1 values ranging between 10-100 ng/mL at 5 ng/mL interval to show the number of observations within a given interval in healthy donors and cancer patients. Comparisons between repeated measures were performed with One-way ANOVA (parametric) or mixed-effects analysis (when values were missing), Friedman test (non-parametric) or Two-way ANOVA with Sidak’s multiple comparisons test. Group analysis between two treatments was performed with two-sided (un)paired student’s *t*-test (parametric) or Mann-Whitney U test and Wilcoxon matched-pairs signed rank test (non-parametric). Univariate and multivariate survival analyses were performed using Cox’s proportional hazards regression model and *P*-values from likelihood ratio tests were reported. *P*<0.05 was considered statistically significant. Statistical analyses were performed with Prism 9 (GraphPad, RRID:SCR_002798) or JMP Pro (v17.0.0).

### Data availability

The data generated in this study are available upon request from the corresponding author.

## Results

### A dominant ∼200 kDa form of sClever-1 is enriched in blood of cancer patients

To elucidate the presence of secreted Clever-1 in biological fluids, we developed a specific Time-resolved fluorescence immunoassay (TRFIA) using Bexmarilimab (Bex) and 9-11 anti-Clever-1 antibodies, optimized for robust sensitivity and stability across different sample material (Figure S1A-E). Next, we analyzed sClever-1 levels in the plasma of healthy donors (n = 32 samples from 21 donors across 1 to 3 timepoints, EDTA blood), in patients with treatment-naïve breast cancer (n = 138, EDTA blood, Table S1), and in patients with various advanced cancers (n = 193, pre-treatment, Li-He blood) participating in the MATINS trial (**Figure 1A**). We observed a highly significant enrichment of sClever-1 in the plasma of patients with cancer compared to undiagnosed healthy donors (**Figure 1B**). In Receiver Operating Characteristic (ROC) analysis the enrichment showed excellent >0.8 performance discriminating cancer patients from healthy donors with 100% specificity for both cohorts at 48.7 ng/mL (**Figure 1C**).

**Figure 1.**
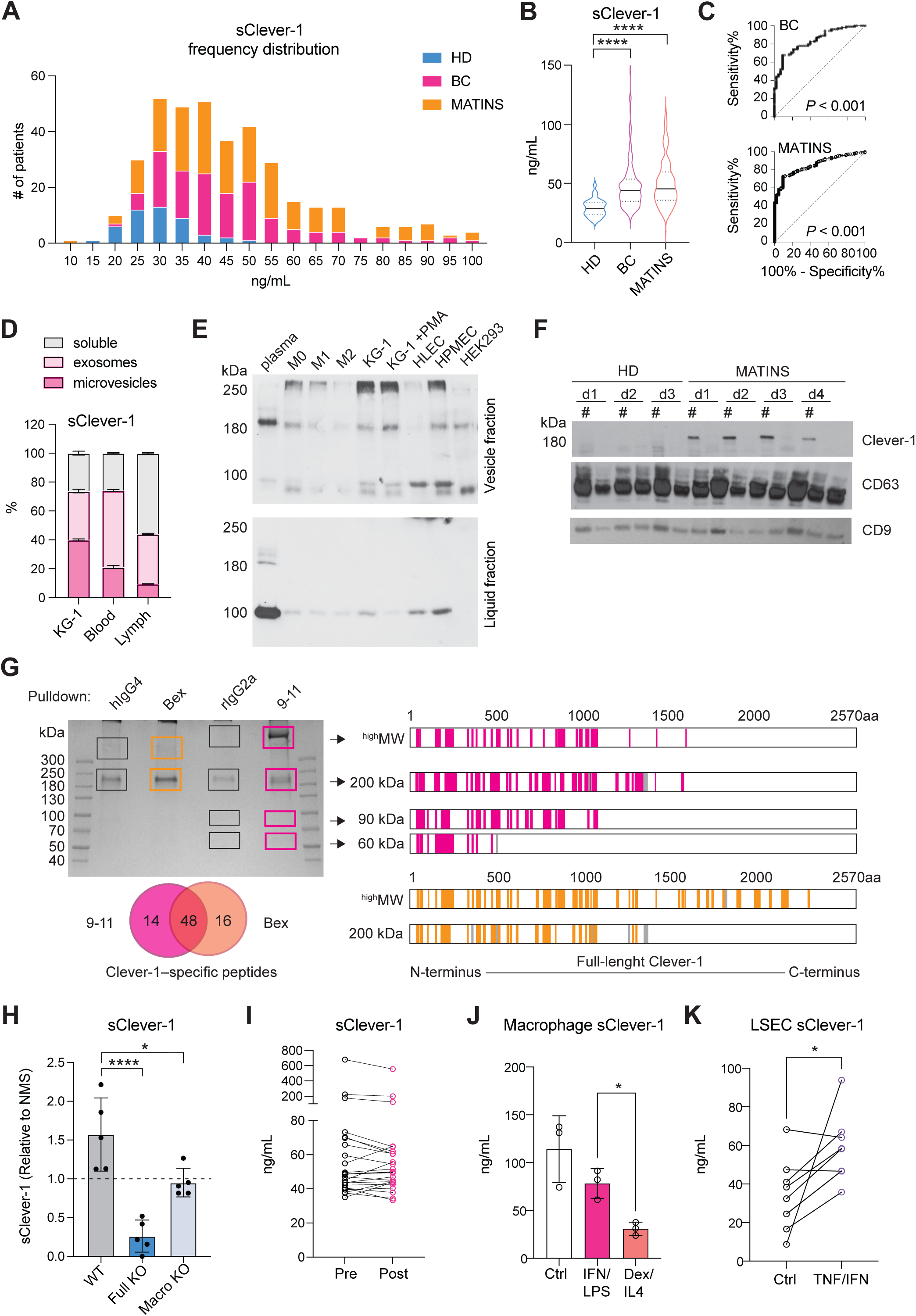
Circulating soluble forms of Clever-1 are enriched in cancer patients. **A**, Frequency distribution and **B**, median of sClever-1 concentration in plasma of healthy donors (HD) (n = 32, blue), patients with treatment naïve breast cancer (BC) (n = 138, magenta) and patients with various advanced solid tumors participating in the MATINS trial (n = 193, orange). One-way ANOVA with Dunnett’s multiple comparison test. **C**, ROC-curves of plasma sClever-1 concentration in BC and MATINS patients. AUC for sClever-1 was 0.84 (CI 0.77-0.90) in BC and 0.85 (CI 0.80-0.90) in MATINS. Specificity for both cohorts at 48.7 ng/mL was 100%. **D**, Quantification (ELISA) of sClever-1 in vesicle fractions in KG-1 cell culture supernatant, blood and lymph separated by ultracentrifugation. Samples were normalized to 1 µg/µL of protein. The graphs represent one independent assay with three technical replicates. **E**, Western blot analysis of sClever-1 detected with 3-372 mouse anti-human Clever-1 antibody (parent of Bex, RRID:AB_3661737) in vesicle and liquid fractions of plasma and cell culture supernatants of primary human macrophages (M0, no polarization; M1, IFNγ + LPS; M2, IL-4 + dexamethasone), endothelial cells (HLEC, human lymphatic endothelial cells; HPMEC, human pulmonary microvascular endothelial cells) and cell lines (KG-1 with/without PMA and HEK293). Equal loading of 10 μg of protein sample per lane was ensured by Qubit measurement of protein concentration. **F**, Western blot showing the abundance of a ∼200 kDa sClever-1 species in healthy donors (n = 3) and MATINS patients (n = 4). # denotes the vesicle fraction. CD63 was used as a loading control. **G**, Mass spec analysis of Coomassie stained bands immunoprecipitated from human serum with Bex or 9-11 anti-Clever-1 antibodies and relative isotype controls hIgG4 and rIgG2a, respectively. The cartoons show Clever-1–specific peptide hits mapped to the full-length Clever-1 protein for Bex (orange) and 9-11 (magenta) pulldown. Venn diagrams depict shared peptides identified by Bex and 9-11. **H**, Quantification (ELISA) of sClever-1 in mouse serum of tumor-bearing wild type (WT, n = 5), full Clever-1 knock-out (Full KO, n = 4) and macrophage specific Clever-1 knock-out (Macro KO, n = 5) mice. The signal was normalized to a pool of normal mouse serum. **I**, sClever-1 concentration in plasma of breast cancer patients before (pre) and three weeks after (post) mastectomy (n = 24). **J**, sClever-1 levels in macrophages stimulated with IFN/LPS or dexamethasone/IL4. Repeated measures One-way ANOVA (n = 3). **K**, sClever-1 levels in primary human liver sinusoidal endothelial cells (LSEC) treated with or without TNFα and IFNγ. Paired Student’s t-test (n = 8). * *P* <0.05, **** *P* <0.0001.

Since healthy controls had significantly lower age than the patients with breast cancer (ΔAge: 27 years, *P*<0.0001), we analyzed whether age correlated with plasma sClever-1 levels. As we found a weak, but significant correlation between ln-transformed sClever-1 and age in breast cancer (Figure S2A), we next performed multiple linear regression to evaluate the effects of age and cancer on plasma sClever levels (Figure S2B). While both age and the presence of cancer were significant predictors of plasma sClever-1, our model predicted the presence of cancer to increase sClever-1 levels as much as 60 years of aging (Table S2, Figure S2C). In contrast to treatment-naïve breast cancer patients, we did not observe an overall correlation between age and sClever-1 in the MATINS trial patients, which represented ten different advanced cancer cohorts and had variation in previous lines of therapies (Figure S2A, D and E). Male gender was associated with slightly higher sClever-1 levels in both healthy donors and the MATINS trial patients. However, in regression analyses gender did not significantly explain sClever-1 levels and gender-related differences in sClever-1 levels would only underestimate rather than overestimate the observed effect of cancer on sClever-1 levels in the breast cancer cohort (data not shown).

To characterize the secretory profile of sClever-1 for its distribution between extracellular vesicles and a freely soluble form, we fractionated the culture supernatant from Clever-1-expressing acute myeloid leukemia cells (KG-1), human blood and lymph by ultracentrifugation. This approach enabled us to determine sClever-1 levels in fractions containing microvesicles, exosomes and soluble proteins. All the studied fractions showed detectable sClever-1 protein in which lymph had the highest content of sClever-1 in the soluble form (**Figure 1D**). We further assessed the release of sClever-1 in the vesicle versus soluble fraction from various primary human cells and cell lines by Western blot to identify the cell type producing sClever-1 into blood. Different from the full-length ∼280 kDa Clever-1 band, the plasma contained a dominant and abundant ∼200 kDa sClever-1 species, which was observed in all macrophage subsets and endothelial cells as well as PMA-differentiated KG-1 cells (**Figure 1E**). Direct comparison of healthy plasma with MATINS patient’s pre-treatment plasma in the vesicle fraction showed increased abundance of the ∼200 kDa sClever-1 (**Figure 1F**).

For validating the Western blot data for sClever-1 we performed mass spec analysis of each gel band containing serum protein immunoprecipitated via biotinylated anti-Clever-1 antibody (Bex and 9-11) (**Figure 1G**). The analysis revealed peptide sequences from full-length Clever-1 isoform #1. Further, whilst there was an abundance of peptides unique to full-length Clever-1 isoform #1, there were no peptides unique to the predicted isoform #2. This was true even for the 60 kDa and 90 kDa bands immunoprecipitated via 9-11 antibody. Thus, it can be deduced that the hypothetical secreted isoform #2 variant of Clever-1 was not obtainable as a protein from serum. An interesting observation from the mass spec analysis was that each species of sClever-1 that was smaller than full-length Clever-1 was heavily truncated at the C-terminus. A confirmatory mass spec was performed from serum where whole pulldowns were directly analyzed without gel excision. The analyses showed an overwhelming peptide coverage for the first half of Clever-1 and little beyond the end of the half-molecule, confirming that the **∼**200 kDa species of sClever-1 from serum is the dominant one, and that a recombinant secreted first half of Clever-1 protein would be a good approximation of sClever-1 (data not shown).

To further investigate whether the tumor microenvironment was a source of sClever-1 and how the different cell types contributed to the secretion of sClever-1 we modified our ELISA assay to detect mouse Clever-1 and used that to measure sClever-1 in Clever-1 full knock-out (KO) or macrophage-specific Clever-1 KO mice described in our previous study (25). Compared to pooled normal mouse serum, tumor-bearing wild type mice showed an increase in sClever-1 (**Figure 1H**). As expected, the tumor-bearing full KO mice showed a significant drop in sClever-1, which was not as dramatic in the tumor-bearing macrophage KO mice suggesting that the endothelial cells contribute half to the observed sClever-1 in murine blood (**Figure 1H**). We also measured sClever-1 in the plasma of the breast cancer patients three weeks after their surgery and compared that to their pre-surgical blood sample. Overall sClever-1 dropped ∼10% after tumor removal but was not significantly reduced (**Figure 1I**), indicating a more systemic source of sClever-1 even though exposure of human primary macrophages to breast cancer cell conditioned medium upregulated sClever-1 (Figure S3). Importantly, chronic inflammation is reported to induce Clever-1 expression on endothelial cells (26) and Clever-1 expression on monocytes is rapidly decreased during pre-eclampsia (4). Thus, we quantified sClever-1 release in macrophage stimulated with IFNγ + LPS or dexamethasone + IL4 (**Figure 1J**). Interestingly, IFNγ/LPS-stimulated macrophages released more sClever-1 than the ones stimulated with immunosuppressive factors, and was not due to shedding of Clever-1 from the cell surface as the level remained the same or even increased after IFNγ/LPS (Figure S3). Similarly, primary human liver sinusoidal endothelial cells (LSEC), which have high expression of Clever-1, adapted to an inflammatory stimulus by increasing the release of sClever-1 (**Figure 1K**).

### Recombinant production and quality control of the ∼200 kDa sClever-1

To investigate the function of the cancer enriched form of sClever-1 we recombinantly produced a half Clever-1 molecule (H1) mimicking the ∼200 kDa sClever-1 (Figure S4A). The amino acid sequence of H1 consists of the first half of full-length Clever-1 and encompasses the peptide sequences identified by the pulldown and mass spec analyses of sClever-1 from plasma, and has a predicted MW of approximately the same size as sClever-1. The secreted protein was subsequently purified using Ni^2+^-affinity and size-exclusion chromatography to >95% purity (Figure S4B). H1 elutes as a broad, homogeneous monomeric peak with a minor, highly oligomeric fraction present (Figure S4C). The absolute MW of H1 agreed well with the sequence-derived molar mass and further suggested the presence of 18 ± 2% glycosylation by mass. The purified protein was properly folded with a similar secondary structure composition as predicted by AlphaFold2 (Figure S4D). H1 was found to be stable and well-behaved (Figure S4E), even after several rounds of freeze-thaw cycles (data not shown). The broad elution profile of H1 can be explained by the glycan content and a high order of inter-domain flexibility as assessed by dynamic light scattering (DLS) (Figure S4F), size-exclusion chromatography and small-angle X-ray scattering (SEC-SAXS) (Figure S4G).

### H1 binds activated T cells at the same time as transient phosphatidylserine exposure during early T cell activation

Having performed the quality control for H1 we investigated its binding on peripheral T cells enriched by CD3 negative selection. In non-activated T cells positive binding was observed in a FCS^low^ population most likely representing apoptotic cells (**Figure 2A**). Indeed, previous reports indicate that Clever-1 mediates PS-dependent clearance of cell corpses in alternatively activated macrophages (27). However, when the lymphocytes were activated via T cell receptor (TCR) engagement a FSC^high^ population became H1 positive, which was more prominent than H1 binding to the FSC^low^ population in both non-activated and activated states after 48h (**Figure 2B**). To verify that the FCS^high^ cells were not apoptotic we stained them with Annexin V and FAM-VAD-FMK, which is a caspase inhibitor that blocks apoptosis and because of its carboxyfluorescein can be used to detect early apoptotic cells. As expected, the FSC^low^ H1^+^ cells produced a high signal for both markers, which was not observed in the FSC^high^ H1^+^ cells (**Figure 2C**). Pre-incubation of H1 with the anti-Clever-1 antibodies, bexmarilimab or 9-11, did not block H1 binding to T cells (Figure S5A), suggesting that the T cell interaction was occurring via a non-competing epitope. Due to the previously identified possible interaction of Clever-1 with PS (27) we pre-incubated the activated T cells with Annexin V to prevent H1 binding on the cells. This had no effect on H1 binding (Figure S5B) indicating that H1 was binding T cells via another ligand exposed during phospholipid translocation (flipping) from one bilayer to the other.

**Figure 2.**
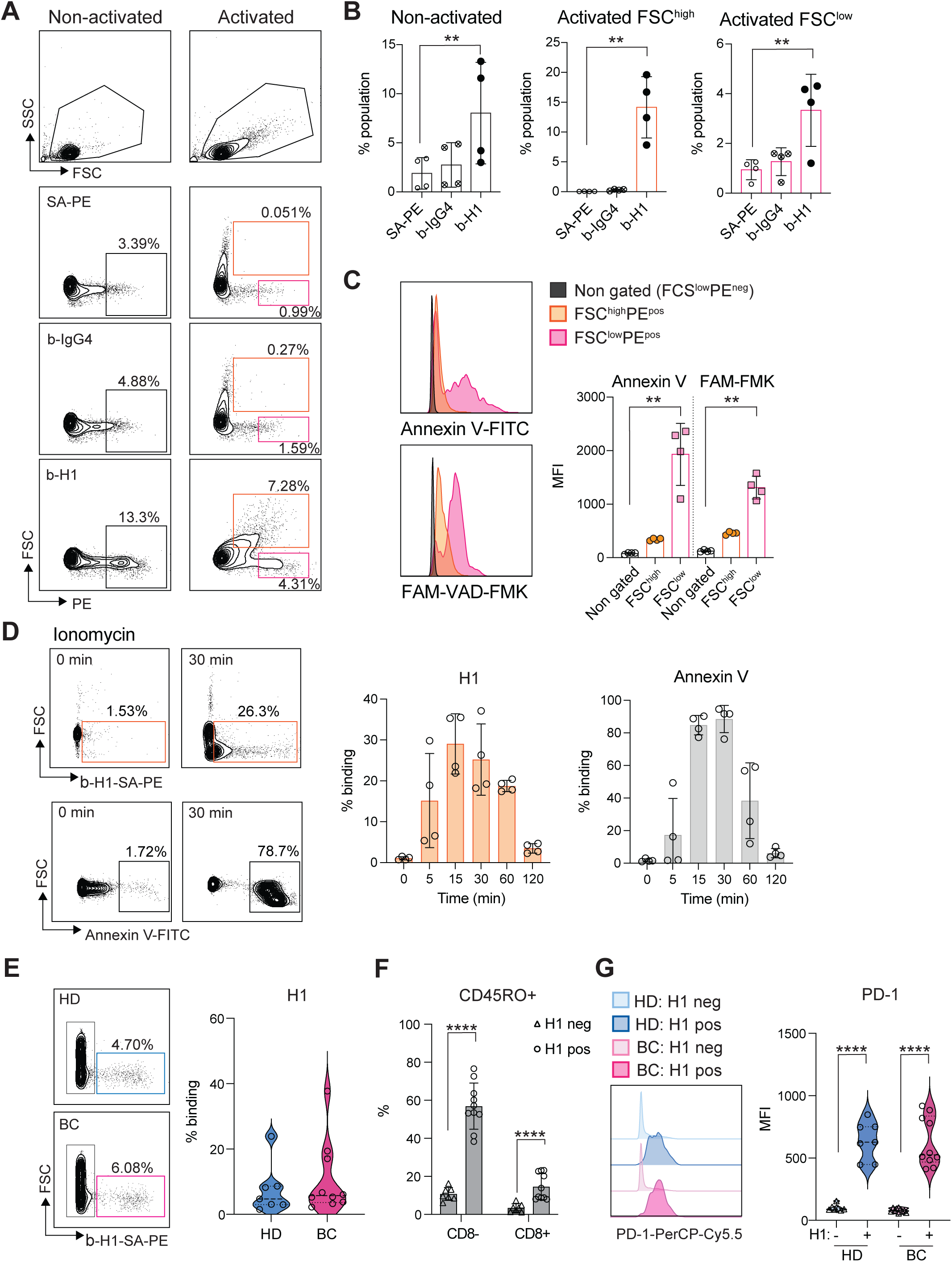
Clever-1 binds activated lymphocytes during transient phosphatidylserine exposure. **A**, Flow cytometry plots **B**, and quantification of H1 binding on primary T cells after 48 hours of activation with anti-CD3/CD28 and IL2 (FCS^high^ gate, orange; n = 4 donors). Streptavidin-phycoerythrin (SA-PE) and biotinylated human IgG4 served as negative controls for H1 staining. **C**, Co-staining of activated T cells with Annexin V (to detect cell surface phosphatidyl serine) and FAM-VAD-FMK (to detect pre-apoptotic events) showing apoptotic cells in the FCS^low^ gate. MFI, median fluorescence intensity. One-way ANOVA with Dunnett’s multiple comparison test (B and C). **D**, H1 and Annexin V binding kinetics on primary human T cells activated by ionomycin. **E**, Flow cytometry analysis of H1 binding on lymphocytes obtained from healthy donors (HD, n = 7) and breast cancer patients (BC, n = 10). **F**, Analysis of CD45RO expression in CD8- and CD8+ cells binding H1 in comparison to H1 negative cells. Paired Student’s t-test. **G**, Expression of PD-1 in H1 positive and negative cells from HD and BC patients. Paired Student’s t-test. * *P* <0.05, ** *P* <0.01, *** *P* <0.001, **** *P* <0.0001.

Externalization of PS is not restricted to apoptotic cells and can occur upon cellular activation in a transient manner where elevated cytoplasmic Ca^2+^ followed by induction of phospholipid scramblase accelerates phospholipid movement (11). As ionomycin is known to increase intracellular Ca^2+^ levels, we tested whether short term activation of T cells with ionomycin can increase H1 binding. Already at five minutes H1 binding increased and peaked around 30 minutes resembling similar kinetics as seen with Annexin V (**Figure 2D**).

To get insight on the cell populations possibly affected by sClever-1 in blood we analysed H1 binding on peripheral lymphocytes collected from healthy donors and breast cancer patients. H1 binding was observed to be similar between healthy donors (median 4.7 range 22.4%) and breast cancer patients (median 5.7 range 35.5%) (**Figure 2E**), where H1 positive signal was mostly observed on CD45RO^+^ memory cells in both CD8 positive and negative cell populations (**Figure 2F** and Figure S5C). The H1^+^ cells had significantly higher expression of PD-1 compared to cells not binding H1 indicative of selective binding of sClever-1 on activated memory T cells (**Figure 2G**).

### H1 disrupts TCR activation and leads to impaired Th1 expansion

We next investigated whether H1 binding on T cells has functional consequences during their activation and proliferation. In the presence of H1 the CD8^+^ T cells reached four generations during the six-day incubation period. Hence, no dramatic effect on proliferation compared to control treated cells was observed (**Figure 3A**). When looking at their differentiation based on lineage determining transcription factors T-bet and FoxP3, the H1 treated cells had a significantly higher percentage of FoxP3^+^ cells compared to control cells (**Figure 3B**). Since suboptimal TCR activation has been reported to increase upregulation of FoxP3 on CD8^+^ T cells (28) we assessed the early signalling events downstream of TCR. By binding to T cells H1 was able to impair the phosphorylation of Lck at Y394 (**Figure 3C**). Similarly, in a Jurkat NFκB reporter cell line, the presence of H1 tended (*P* = 0.091) to reduce reporter activity (**Figure 3D**). FoxP3 is known to suppress the function of NFAT and NFκB, and this leads to suppression of the expression of many genes including IL2 and effector T cell cytokines (29). Indeed, cytokine profiling of primary T cell cultures showed a significant reduction in IL1β, IL6, IL10 and IL17 among the measured 27 cytokines when incubated with H1 (**Figure 3E**). These cytokines were not directly indicative of inducing any specific lineage differentiation of the activated T cells, thus we functionally tested how the T cells polarized autologous primary human macrophages. The H1 cultured T cells promoted a polarization change in macrophages seen as a significant downregulation of HLA-DR and CD40, whereas Clever-1 expression on macrophages was upregulated suggesting a more immunosuppressive phenotype induced by the H1-treated T cells (**Figure 3F**).

**Figure 3.**
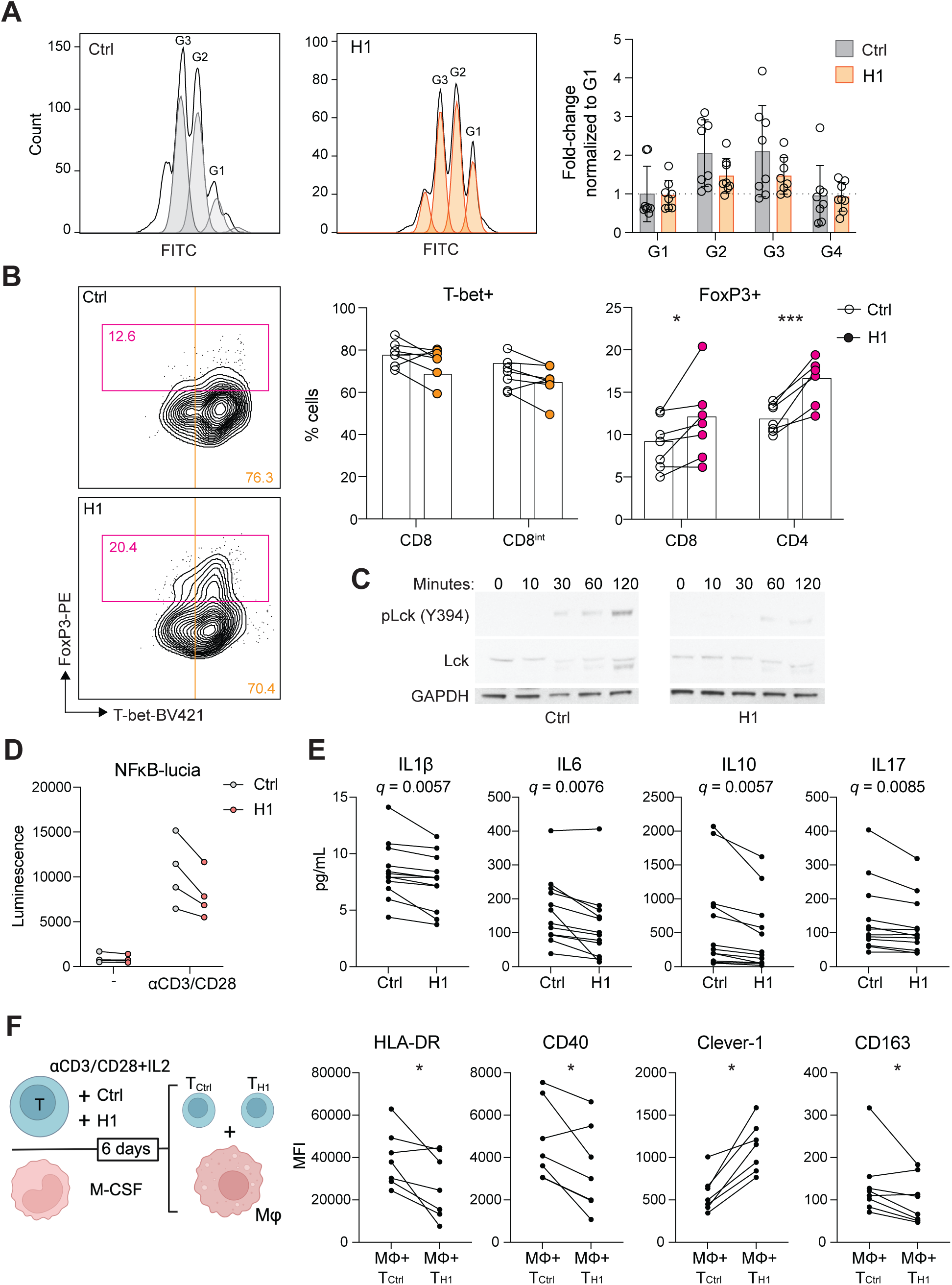
Clever-1 promotes the differentiation of suppressive FoxP3^+^CD8^+^ T cells. **A**, Proliferation of CD8^+^ T cells after six days in culture with H1 (50 µg/mL) as assessed by CFSE dilution and quantification of cell numbers relative to control (BSA) treated cells (n = 8 donors). **B**, Flow cytometry plots showing CD8^+^ populations from which T cell lineage markers T-bet and FoxP3 were quantified (n = 7 donors). Paired t-test. **C**, Representative Western blot of Lck phosphorylation at different time points after T cell activation with or without H1. Representative blots from four independent experiments. **D**, Jurkat NFκB reporter activity induced after TCR-ligation with or without H1. Graph showing four independent experiments performed in triplicates. **E**, Cytokine analysis of primary T cell cultures incubated with or without H1 (n = 10 donors). Significance shown as FDR-adjusted q-values for multiple t-tests. **F**, Schematic of the assay design in which primary T cells were activated with or without H1 for six days and in parallel monocytes from the same donors were differentiated into macrophages. Thereafter, the T cells were washed and incubated with the macrophages for 48h and polarization markers expressed by macrophages (CD64^+^) were assessed with flow cytometry (n = 6 donors). Paired t-test. * *P* <0.05, *** *P* <0.001.

### H1 binds IGF2R on T cells via mannose-6-phosphate

Several ligands have been identified to bind Clever-1 such as acLDL, SPARC and chitinase like proteins and recently serum periostin, reelin, and TGFBi (30). To identify the counter receptor for sClever-1 on T cells we first screened H1 binding on a HEK293 cell-based Retrogenix platform consisting of 6449 human plasma membrane proteins, tethered human secreted proteins and 397 heterodimer plasma membrane protein complexes. Testing H1 binding on fixed cells by two different staining protocols did not yield any hits. Thus, we next tested H1 binding on Jurkat cells to utilize them in affinity pulldown assays. To our surprise, H1 showed very good binding to all Jurkat cells even without prior TCR-stimulation. The binding was not induced by apoptosis since puromycin treated Jurkat cells did not bind H1 (**Figure 4A**). Mass spec analysis of proteins pulled down with H1 revealed several hits involved in T cell activation such as IGF2R, TFRC, Zap70, CCT8, CCT6A (**Figure 4B**).

**Figure 4.**
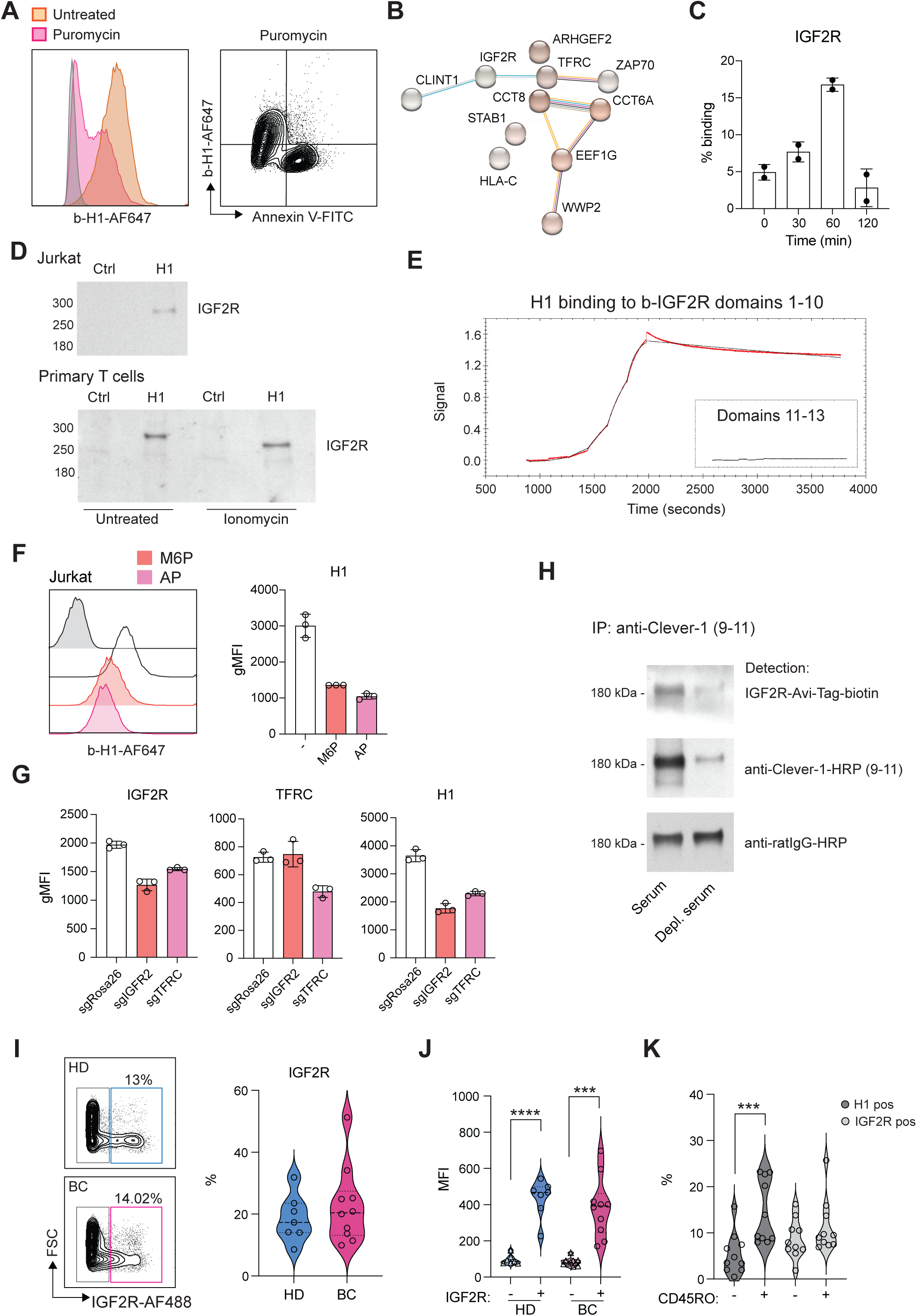
Clever-1 binds to IGF2R via mannose-6-phosphate. **A,** Flow cytometry plots showing H1 and Annexin V binding on Jurkat cells after apoptosis induction (5 μg/mL of puromycin for 3 hours). **B,** STRING database network summary of mass spectrometry proteomics results from pulldowns with biotinylated-H1 from Jurkat cell membranes. **C,** Flow cytometry analysis of cell surface IGF2R expression on ionomycin stimulated primary human T cells (n = 2 donors). Experiment repeated twice. **D,** Western blots of pulldowns with biotinylated-H1 detected with anti-IGF2R. The primary T cells were stimulated with ionomycin for 30 minutes before performing the pulldowns. Control samples contained beads only. **E,** Octet BLI analysis of the binding of H1 to IGF2R domains 1-10 and 11-13. Single cycle kinetics data was fitted using a 1:1 Langmuir binding model with reference subtraction, to estimate the binding affinity. **F,** Flow cytometry of H1 binding to Jurkat cells in the presence of excess mannose-6-phosphate (M6P) or after alkaline phosphatase (AP) treatment. **G,** Flow cytometry of CRISPR/Cas9 edited Jurkat cells for H1 binding. gRNA against Rosa26 was included as a negative control. **H,** Western blot to detect M6P on endogenous serum sClever-1 with recombinant IGF2R-His-AviTag protein. Biotinylated 9-11 antibody was used to pulldown sClever-1 from either human AB serum, or human AB serum depleted of sClever-1 by pulldown with 9-11 (confirmed via ELISA). Equal loading was confirmed with an anti-rat secondary antibody. Note, due to the non-reducing conditions of the gel, any streptavidin that leached from the Dynabeads would remain in complex with the biotinylated 9-11 antibody, creating a band larger than the anticipated 150 kDa for an IgG2a. **I**, Flow cytometry analysis of IGF2R expression on lymphocytes obtained from healthy donors (HD, n = 7) and breast cancer patients (BC, n = 10). **J**, Expression of PD-1 in IGF2R positive and negative cells from HD and BC patients. Paired Student’s t-test. **K**, Percent of CD45RO positive cells binding H1 or expressing IGF2R in relation to CD45RO negative cells. Paired Student’s t-test. *** *P* <0.001, **** *P* <0.0001.

We have previously identified an interaction of cellular Clever-1 with transferrin receptor 1 (TFRC) in primary human macrophages (5). Thus, we decided to investigate IGF2R in more detail as it was not included in the Retrogenix assay nor would endogenous IGF2R normally be displayed upon the cell surface of HEK293 cells. We observed a similar upregulation of IGF2R on the membrane of primary human T cells after ionomycin stimulation as we observed for Annexin V in primary human T cells (**Figure 4C**). Indeed, it has previously been published that an increase in cell surface IGF2R is a transient and very early marker of primary T cell activation (31). Pulldown assays with Jurkat and primary human T cells confirmed the interaction of H1 with IGF2R (**Figure 4D**). Using recombinant IGF2R in Octet we were able to validate that H1 bound directly to IGF2R in domains 1-10 but not in domains 11-13 known to bind IGF II (**Figure 4E**). Since IGF2R is a cation-independent mannose-6-phosphate receptor we tested whether an excess amount of M6P or removing M6P from H1 by alkaline phosphate treatment would abolish H1 binding to IGF2R. On Octet, H1 binding to IGF2R was completely abolished with M6P manipulation (data not shown). Also, on Jurkat cells H1 binding was decreased with either an excess amount of free M6P or removal of M6P from H1 (**Figure 4F**). To fully demonstrate H1 binding on T cells via IGF2R, we knocked out either IGF2R or TFRC from Jurkat cells by CRISPR/Cas9. We were able to achieve a ∼30% reduction in the expression levels of both proteins and this was able to decrease H1 binding by ∼50% to IGF2R CRISPR cells (**Figure 4G**). It is interesting to note that gene silencing of TFRC downregulated IGF2R and therefore reduced also binding of H1 to TFRC CRISPR cells (**Figure 4G**). To confirm the interaction of endogenous sClever-1 with IGF2R, we performed Clever-1 pulldowns from serum and used a recombinant IGF2R containing an AviTag as detection reagent. Positive signal was observed with IGF2R when sClever-1 was detected, and was clearly reduced when the pulldowns were performed with Clever-1 depleted serum (**Figure 4H**). In the circulation, IGF2R expressing lymphocytes were detected both in healthy donors (mean 18.6 ±7.6%) and breast cancer patients (mean 22.7 ±12.4) with high PD-1 expression (**Figure 4I-J** and Figure S5C). Unlike H1 binding on CD45RO^+^ memory cells, a similar preference was not observed for IGF2R expression (**Figure 4K**) proposing a broader expression profile for IGF2R on activated T cells.

### Bexmarilimab treatment inhibits the release of sClever-1

To understand the contribution of endogenous sClever-1 in regulating T cell tolerance we generated vesicle fractions from conditioned media of primary human macrophages silenced for Clever-1 (**Figure 5A**) and tested their effect in a Jurkat-Raji cell assay where antigen specific T cell activation is inhibited by overexpression of PD-1/L1 (**Figure 5B**). This inhibition can be released by anti-PD1 (e.g. nivolumab) leading to activation of the NFAT-reporter. The gene silencing of *STAB1* with sg1 led to a complete depletion of the Clever-1 protein in both cell lysates and the vesicle fraction, whereas the gene silencing of *STAB1* with sg2 was less robust (**Figure 5A**). Interestingly, in Clever-1 CRISPR macrophages CD206 expression was also substantially reduced. Different from gene silencing, the Clever-1-binding antibodies 9-11 and bexmarilimab did not induce downregulation of Clever-1 in the cells but were able to reduce sClever-1 compared to isotype-control-treated macrophages (**Figure 5A**). Notably, while Clever-1 was not expressed by M1 polarized macrophages (IFNγ and LPS), the extracellular vesicles (EV) released from these cells during polarization showed abundant levels of Clever-1. Hence, EVs derived from both M1 and M2 polarized macrophages reduced nivolumab-induced Jurkat reporter activity. In contrast, EVs derived from Clever-1 silenced M2 polarized macrophages were less suppressive (**Figure 5C**). Of note, Clever-1 silencing nor bexmarilimab treatment per se did not contribute to the number of EVs secreted from macrophages (data not shown). Clever-1 itself was exposed on the EV surface since proteinase K treatment removed Clever-1 from the EV fraction detected by Western blot (Figure S6).

**Figure 5.**
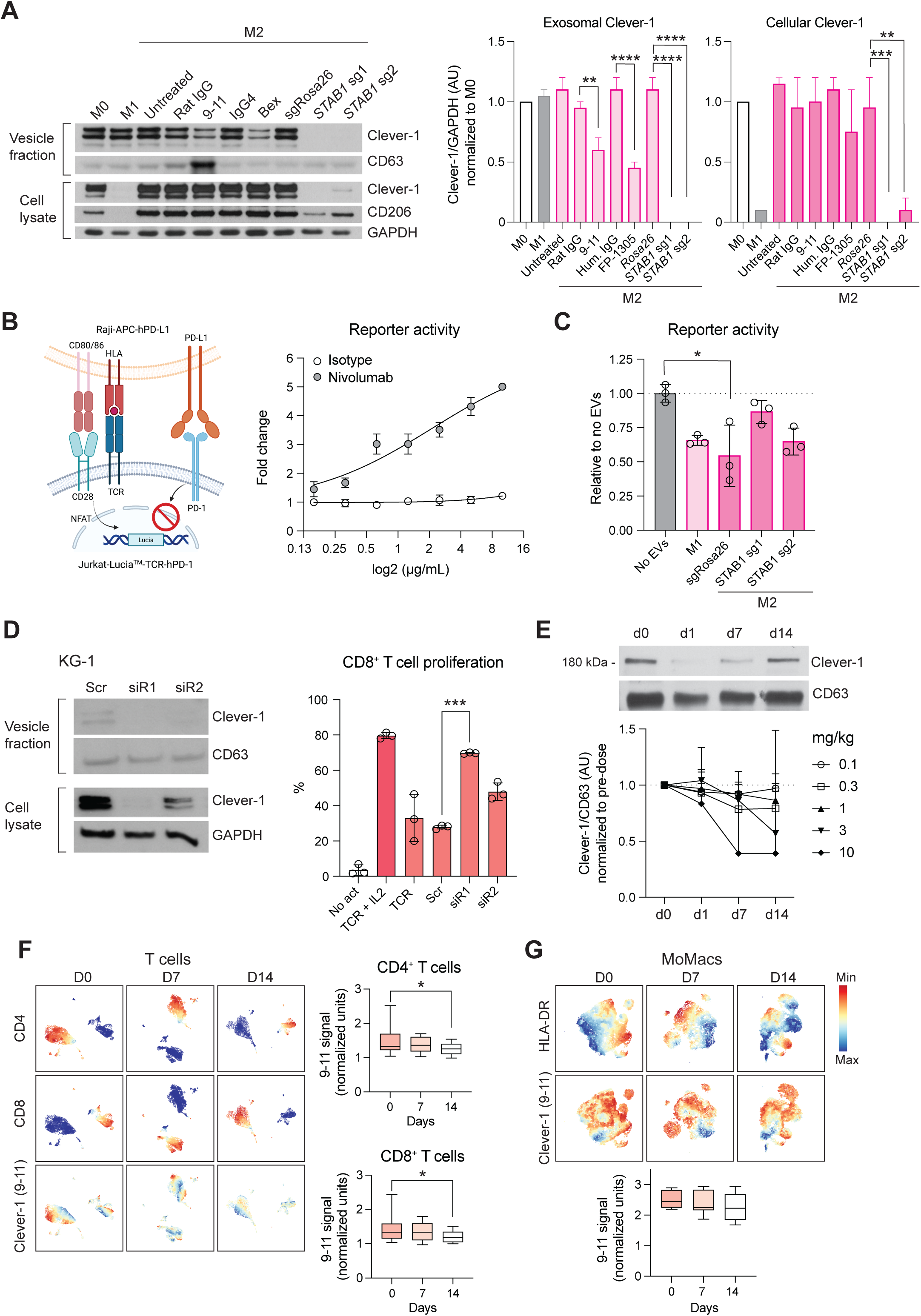
Bexmarilimab inhibits the release of sClever-1. **A,** Representative Western blot and quantification of Clever-1 levels in cells and the secreted vesicle fraction after anti-Clever-1 treatment with 9-11 or Bex, or Clever-1 silencing by CRISPR/Cas9 (*STAB1* sg1 and sg2) compared to cells treated with isotype or non-targeting guide RNA (sgRosa26) control, respectively. CD63 was used as loading control and CD206 for M2 polarization control. Clever-1 signal was detected with anti-Clever-1 (4G9) antibody. GAPDH loading control was used for normalizing signal intensity for Clever-1. M0 signal was considered as 1. Student’s t-test (n = 3). **B**, Schematic of the Jurkat-Raji cell assay and representative reporter activity at different doses of nivolumab in comparison to isotype control treatment. **C**, Nivolumab-activated Jurkat-Raji cell (no EVs) reporter activity in the presence of EVs collected from primary human macrophages shown in A. Kruskal-Wallis test with Dunn’s multiple comparison. **D**, Western blot analysis of Clever-1 after siRNA knock-down in KG-1 cells and the effect of EVs (2.5 μg/mL) collected from these cells in regulating CD8^+^ T cell proliferation. Paired t-test. **E**, Representative Western blot of the ∼200 kDa sClever-1 band from the plasma vesicle fraction of a patient administered with 0.1 mg/kg of bexmarilimab and quantification at different dose levels (0.1-10 mg/kg, n = 3 in all doses except n = 2 at 10 mg/kg) normalized to CD63 levels. **F**, Hierarchical Stochastical Neighbor Embedding (hSNE) clusters of mass cytometry data of CD4^+^ and CD8^+^ T cells and heatmaps of anti-Clever-1 (9-11 antibody) immunoreactivity during the first treatment cycle with bexmarilimab. Bexmarilimab administration shows significantly decreased binding of 9-11 antibody on CD4^+^ and CD8^+^ T cells on day 15 indicating that less sClever-1 is bound to lymphocytes. **G**, The changes of membrane bound Clever-1 levels in monocytes/macrophages (MoMacs) are shown as reference. Mixed-effects analysis with Holm-Sidak’s multiple comparison test, n = 6-7. * *P* < 0.05, ** *P* < 0.01, *** *P* < 0.001, **** *P* < 0.0001.

Since bexmarilimab is also clinically evaluated for the treatment of acute myeloid leukemia we silenced Clever-1 from the KG-1 AML cell line and incubated isolated EVs with activated primary CD8^+^ T cells. The EVs from control treated cells did not suppress CD8^+^ T cell proliferation (**Figure 5D**). However, when EVs from Clever-1 siRNA silenced KG-1 cells were applied a significant induction of proliferation was observed reaching the level of proliferation of cells supplemented with IL2.

We further studied the effect of bexmarilimab in reducing the release of sClever-1 in cancer patients by analysing the plasma vesicle fractions of patients participating in the MATINS trial. The patients were dosed with 0.1-10 mg/kg of bexmarilimab and the level of cancer-enriched ∼200 kDa sClever-1 was analyzed across two weeks post treatment. By day 14 sClever-1 was reduced with the most reduction observed with higher doses, although variation between patients occurred (**Figure 5E**). When we looked at the MATINS patient blood samples for sClever-1 binding on T lymphocytes at pre-dose and 1-2 weeks after treatment we observed that a subset of CD8^+^ T cells, that initially showed positivity for sClever-1, significantly decreased 9-11 (anti-Clever-1) signal by two weeks on treatment (**Figure 5F**). A similar decrease was observed on CD4^+^ T cells but not on monocytes/macrophages where 9-11 binds to full-length Clever-1 expressed by these cells (**Figure 5G**). This suggests that administration of bexmarilimab can reduce sClever-1 levels in cancer patients and thereby decrease sClever-1 engagement with T cells potentially contributing to the peripheral T cell activation observed in the MATINS trial patients (5). Altogether, these data demonstrate that macrophage-derived Clever-1 containing EVs contribute to T cell suppression and anti-PD-1 resistance, which can be inhibited by bexmarilimab treatment.

### Elevated sClever-1 levels are linked to resistance against anti-PD-1 therapy

To investigate whether sClever-1 levels associate with response to immunotherapy, we obtained a dataset of lung carcinoma and melanoma patients with plasma sClever-1 levels measured before start of anti-PD-1 or anti-PD-L1 immunotherapy (Supplementary Table S3.) To evaluate whether sClever-1 associates with overall survival in patients receiving immunotherapy, we first performed univariate Cox proportional hazards regression analyses for each parameter in the dataset and selected those significantly associating with overall survival for a multivariate survival analysis alongside sClever-1. In the multivariate analysis, the number of received immunotherapy doses associated most significantly with overall survival (*P*-value = 0.0002, hazard ratio 0.75 [95% CI: 0.62 – 0.88]), while sClever-1 or parameters commonly improving prognosis (e.g. lymphocyte count, combination with chemotherapy) failed to show significant associations, presumably reflecting to too high inter-patient variation in this small dataset including different immunotherapeutics, doses, dosing intervals, cancer types and treatment procedures. Therefore, we only describe here preliminary associations that should be confirmed in a larger and more homogenous dataset.

We selected the number of received immunotherapy doses as a surrogate marker for anti-PD-(L)1 efficacy, as it most highly correlated with overall survival. Receiving ≤4 doses was considered as no response to treatment and >4 doses was considered as responsive to treatment. This cut-off was selected based on the reported mean time-to-response of 2.2 months for nivolumab in the Checkmate:073 phase III trial (32) considering that the treatment is given Q3W. Also, the trial reported progression-free survival of 3.1 months in patients who did not show benefit, which supports our selected cut-off. We observed higher sClever-1 levels in patients receiving ≤4 anti-PD-1 immunotherapy doses compared to those who received >4 (**Figure 6A-E**). The difference was significant, when looking only at anti-PD-1−treated patients, who had not received chemo- or radiotherapy. Therefore, we analyzed sClever-1 association with overall survival in these patient subsets separately. The analyses suggested higher sClever-1 levels to be a worse prognostic factor for patients receiving anti-PD-1 therapy (**Figure 6F**).

**Figure 6.**
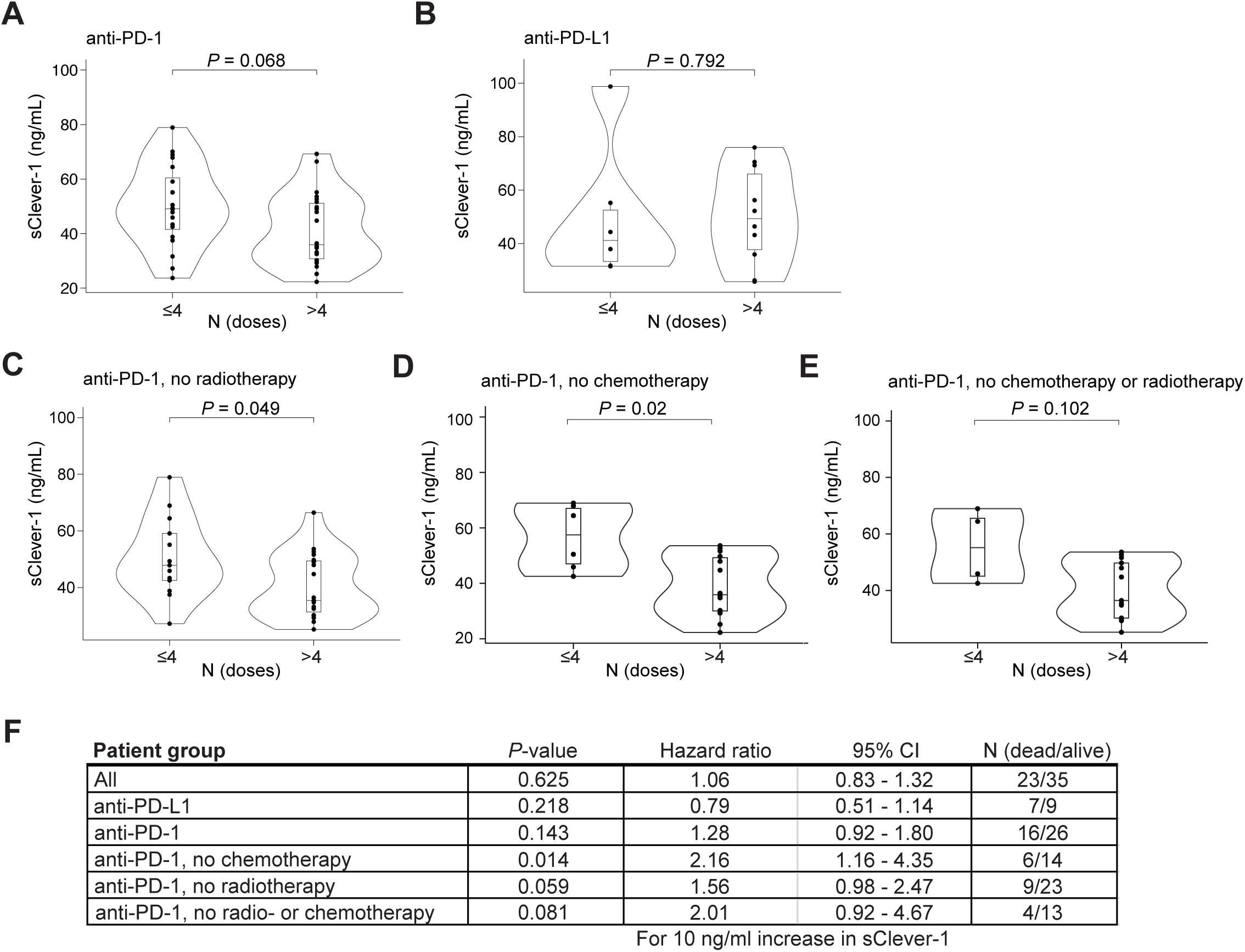
High sClever-1 levels associate with resistance to anti-PD-1 therapy. **A-E,** sClever-1 levels (ng/mL) in patients receiving ≤ or > 4 doses of indicated immunotherapy. Tukey boxplots show median and interquartile range, groups compared with Wilcoxon rank sum test. Analyses were performed in subgroups of patients receiving anti-PD-1 therapy (A), anti-PD-L1 therapy (B) or anti-PD-1 therapy and no chemotherapy (C), radiotherapy (D) or chemo- and radiotherapy (E). Points indicate individual patients. **F**, sClever-1 association with overall survival in patient subgroups presented in A. Results from univariate Cox proportional hazards regression models: P-values from likelihood ratio test and hazard ratios with 95% confidence intervals for 10 ng/mL increase in sClever-1 levels.

## Discussion

Clever-1 is known to govern tolerogenic activities on macrophages but it is not clear if it has direct immunosuppressive properties. Here we report the secretion of a truncated form of Clever-1 by macrophages and endothelial cells that bind T cells causing suboptimal T cell activation and promotes the differentiation of suppressive FoxP3^+^CD8^+^ T cells. We identify secreted Clever-1 to be particularly enriched in the plasma of cancer patients indicating a possible broad systemic immunosuppressive role for this receptor.

The reason for elevated sClever-1 levels in cancer is not entirely clear but may reflect pre-existing chronic inflammation often associated with cancer. Indeed, it has been previously reported that breast cancer patients have alterations in several cytokines (both pro-inflammatory and anti-inflammatory), including IFNγ and TNFα (33). The underlying inflammatory cause for increased sClever-1 is further supported by our data showing the induction of the release by IFNγ/TNFα and the sustained sClever-1 levels after tumor surgery. Thus, the secretion of sClever-1 is most likely not specific to cancer but a reflection of a patient’s inflammatory status even though macrophages exposed to ER^+^ breast cancer cell medium increased secretion of sClever-1. The conditioned medium of T47D cells has been shown to contain IFNγ and TNFα (34) possibly explaining the observed results.

The precise mechanism of the release of sClever-1 is not understood, but might be a result of its known functions in endosomal recycling and lysosomal release of other proteins such as YKL-39, a monocyte-attracting and pro-angiogenic chitinase-like protein (35) or a by-product of lysosomal clearance of its ligands, acLDL and SPARC (36). While full-length Clever-1 is selectively expressed by immunosuppressive macrophage populations the abundance of sClever-1 in M1-derived EVs was surprising. We hypothesize this phenomenon can be a result of inflammation-induced activation of secretory pathways contributing to sClever-1 release. This phenomenon may have two possible biological functions. Firstly, upon M1-polarization, macrophages may need to rapidly eliminate Clever-1 to be able to respond to inflammatory cues. Secondly, the EV-derived sClever-1 may reduce the extent of activation of bystander lymphocytes in the surrounding microenvironment to prevent excessive inflammation. Indeed, several negative feedback mechanisms such as secretion of IL10, IDO1 and TGFβ are induced during immune activation to maintain homeostasis and prevent excessive immune responses that could lead to tissue damage or autoimmune diseases.

As most of the protein was found as a significantly truncated ∼200 kDa form lacking a transmembrane domain, proteolytic activity in the endosomes may cause Clever-1 truncation. At least we did not observe any signs of alternative splicing of the receptor in its secreted form nor shedding of the receptor on the cell surface. Further, during the process of EV formation and release, PS can be externalized to the outer leaflet of the membrane to which sClever-1 might in theory remain bound to in the circulation. Externalization of PS is a well-known feature of apoptotic cells, where it serves as an “eat me” signal, facilitating the recognition and clearance of dying cells by phagocytes. However, Fischer and colleagues, have identified reversible, transient PS exposure to be a very early surface marker of cell activation in T cells (11). Also, PS re-distribution has been shown to regulate non-apoptotic signal transduction in lymphocytes (37).

Annexins are known to bind PS and are highly involved in anti-inflammatory processes, adaptive immunity, modulation of coagulation and fibrinolysis, as well as protective shielding of cells from phagocytosis. We initially hypothesized that sClever-1 can compete with Annexin V from binding to PS on activated T cells. However, this was not the case as we did not observe a difference in H1 binding after Annexin V pre-incubation. Thus, sClever-1 binding was most likely facilitated by another molecular interaction. To identify the counter ligand, we screened thousands of cell surface proteins by the Retrogenix platform using our recombinant H1 protein mimicking sClever-1. Due to the transient nature of the identified interaction of H1 on T cells it is not surprising that transfected and fixed HEK293 cells did not give any hits even though the quality of H1 was confirmed with extensive characterization showing a pure and stable, highly flexible, monomeric glycoprotein. Hence, biophysical and structural characterisation revealed properties of H1 that were not available in the literature and only hypothesized for class H scavenger receptors (38,39).

Thus, we concentrated on Jurkat T cells and performed cell membrane pulldowns with H1. Three of the interaction partners strongly and consistently co-immunoprecipitated with H1 (TFRC, ZAP70, and most abundantly, IGF2R) are associated with the regulation of T cell activity. TFRC has been shown to interact with intracellular ZAP70 and can amplify T cell activation (40). However, we found no direct interaction between H1 and recombinant TFRC, although we were able to measure a direct interaction between H1 and recombinant IGF2R protein on Octet BLI. On the T cell surface, IGF2R expression greatly increases upon T cell activation while CD26 becomes M6P-modified, enabling surface IGF2R to bind to M6P-modified CD26, promoting CD26 internalization and proliferation induced by CD26 co-stimulation (41). Indeed, IGF2R is only typically found upon the surface of early activated T cells (not dormant T cells), *in vitro* and *in vivo*, where free M6P has anti-inflammatory effects (42). IGF2R is vital for early T cell activation because it moves Lck into a favourable position with CD45 on the cell membrane, so that CD45 may activate Lck, which in turn may activate ZAP70 (31). Selective down-regulation of surface IGF2R improves CD8^+^ T cell survival in the context of cell contraction after acute activation (LmOVA bacterial infection), possibly because IGF2R is able to bind and internalize GranzymeB, which is an M6P-modified secreted protease (43). In Tregs, IGF2 enhances regulatory T cell functions via IGF2R stimulation independently of the M6P binding domains of IGF2R (44) In our experiments, H1 led to a decrease in Lck phosphorylation and increased FoxP3 expression and suppressive characteristics, while H1 bound to IGF2R on the surface of activated CD8^+^ T cells and Jurkat cells in a M6P-dependent manner. Given the increased concentration of circulating sClever-1 in the serum of cancer patients, and because, like H1, endogenous sClever-1 protein from AB serum is also somewhat M6P-modified, it is reasonable to hypothesize our findings may represent a contributing factor towards immunosuppression by Clever-1 *in vivo*. According to our data, bexmarilimab does not antagonize sClever-1 binding to T cells, implying that bexmarilimab treatment cannot directly block the immunosuppressive effect of sClever-1. However, our previous work (5) shows that bexmarilimab can alter the trafficking of Clever-1 and impair endosomal acidification, which could lead to the observed reduced levels of sClever-1 on EVs. This is supported by our observations that no consistent changes in overall EV yields were detected after bexmarilimab treatment, as determined by protein concentration measurements. Therefore, we do not believe that bexmarilimab affects EV release. Instead, it likely decreases the generation of an EV-associated soluble form of sClever-1, thereby reducing systemic immunosuppression.

The cell type that is the greatest source of endogenous immunosuppressive sClever-1, or even M6P-modified plasma membrane-anchored full-length Clever-1, is not yet known and will be difficult to determine, given the variety of Clever-1 expressing myeloid and endothelial cell types (for example, liver sinusoidal endothelial cells) (45) and phosphatase activity *in vivo*. We were unable to purify recombinant full length Clever-1 in a way that would allow us to accurately study its potential interaction with IGF2R. However, in our group’s previous publication (5), full length IGF2R was not amongst the proteins that were coimmunoprecipitated from M2 macrophage cell lysates by antibodies against Clever-1. While we observed an increase in sClever-1 levels in plasma of cancer patients with early and late-stage cancer, the tumor itself did not seem to be a major source of sClever-1 based on sClever-1 assessment in plasma of breast cancer patients before and three weeks after mastectomy. However, it is possible to envisage an engagement where an activated T cell may be close enough to a macrophage or endothelial cell to receive a suppressive dose of M6P-modified Clever-1, either alone, or in the context of immunosuppressive EVs. We acknowledge the limited characterization of EVs collected from cells and serum in this study but want to point out three studies where Clever-1 protein can be found in microvesicles and exosomes derived from human monocyte-derived dendritic cells (46), in microparticles derived from endothelial cells (HUVEC) (47) and in serum exosomes (48). Whether or not this is a major contributor to the immunosuppressive functions of Clever-1 *in vivo* remains to be determined. But our discovery of sClever-1 upon EVs in human serum, its positive correlation with cancer and negative correlation with anti-PD-1 response, and its reduced levels after bexmarilimab treatment, provide a promising platform to explore the potential of sClever-1 as a cancer prognostic and therapeutic biomarker.

## Author contributions

SP, MV, ÁÁB and MH designed the study. SP, MV, RS, ÁÁB, JHR, DAP, IB, LT, CRF and MH performed experiments and analyzed data. IK recruited and managed clinical sample collection in the breast cancer cohort. TAS, SS and MH supervised the study. MH wrote the first draft of the manuscript. All authors contributed to the final review.

## Acknowledgements

We wish to thank Teija Kanasuo and Mari Parsama for excellent technical assistance, Blanca Tejeda González and Pieta Mattila for experimental help, the Turku Protein Core, the bioinformatics (J.V. Lehtonen) and structural biology (FINStruct) infrastructure support from Biocenter Finland and CSC IT Center for Science for computational infrastructure support at the Structural Bioinformatics Laboratory, (SBL), ACbo Akademi University. Mass spectrometry analyses were performed at the Turku Proteomics Facility supported by Biocenter Finland. The authors acknowledge MAX IV Laboratory for time on Beamline CoSAXS under Proposal 20220453. Research conducted at MAX IV, a Swedish national user facility, is supported by the Swedish Research Council under contract 2018-07152, the Swedish Governmental Agency for Innovation Systems under contract 2018-04969, and Formas under contract 2019-02496. We acknowledge Fátima Herranz for her help operating the CoSAXS.

