## Supplemental material, figures and tables for "Secreted Clever-1 Modulates T Cell Responses and Impacts Cancer Immunotherapy Efficacy"

### Methods:

#### Time-resolved fluorometric immunoassay (TRFIA) for detecting soluble Clever-1

A TRFIA was generated using two different anti-Clever-1 antibodies (9-11 and biotinylated Bex), and a europium-labelled streptavidin-based detection method to analyze sClever-1 in human samples (Figure S1A). Briefly, Nunc Maxisorp white assay plates were coated with 9-11 (InVivo Biotech) diluted at 10 µg/mL in 0.1M NaHCO3 pH 9.6. The plate was washed six times with 0.1% tween in PBS and blocked with 1% milk powder and 1% gelatin in PBS for 1h at RT. Thereafter, the plasma samples were diluted 1:20 in PBS and incubated for 1h at RT. After six washes, the biotinylated Bex was added for 1h RT. The plate was washed again and Eu-Labeled Streptavidin diluted 1000-fold in assay buffer (Ready-for-use Tris-HCl Buffered NaCl solution) (pH 7.8) was applied to the wells for 30 min at RT. Following the incubation, the plate was washed and developed with Enhancement Solution and measured with Viktor Nivo (all from Perkin Elmer). For measuring mouse sClever-1 from serum the biotinylated Bex was replaced by biotinylated mStab1.26 antibody (InVivo Biotech). Due to the unavailability of a recombinant Clever-1 standard when running the cancer cohorts against healthy controls, the standard curve was produced using a dilution series on human lymph from a healthy donor. To select the best standard for analyzing patient blood, we tested lymph samples from different donors and selected the one where plasma samples diluted 1:20 produced signals in the linear range of the standard curve. To validate that the assay was specific for detecting sClever-1 we depleted sClever-1 from the lymph with anti-Clever-1 (9-11) and generated a standard curve using sClever-1 depleted lymph (Figure S1B). Additionally, to control that the observed signal was not due to the dilution of lymph in PBS, the diminishing amount of lymph was compensated by adding sClever-1 depleted lymph equal to the volume of PBS used for generating each dilution. The standard curve was found to be optimal using dilutions 1:10, 1:15, 1:20, 1:40, 1:80 and 1:160 of human lymph. The observed signal in the TRFIA assay was sClever-1 specific since the maximal observed signal using sClever-1 depleted lymph with 1:10 dilution was 8327 (with lymph $3.3x10^6$). Moreover, lymph diluted with sClever-1 depleted lymph produced an identical standard curve ($R^2$ 0.993) compared to lymph diluted with PBS ($R^2$ 0.993) demonstrating that the observed read-out was not due to dilution of the lymph itself.

By the production of H1 and validation of its binding to bexmarilimab (Figure S1C), the standard curve was replaced by sClever-1 depleted plasma spiked with different concentrations of H1. With the new standard, the absolute levels of sClever-1 in the plasma of patients with cancer and healthy controls were retrospectively calculated by running assay control samples containing low, medium and high sClever-1 with the lymph standard in parallel with the H1 standard (Figure S1D). The stability of sClever-1 was tested and showed that the signal remained stable for at least five freeze-thaw cycles (-70°, n = 7) in the tested matrices (Figure S1E).

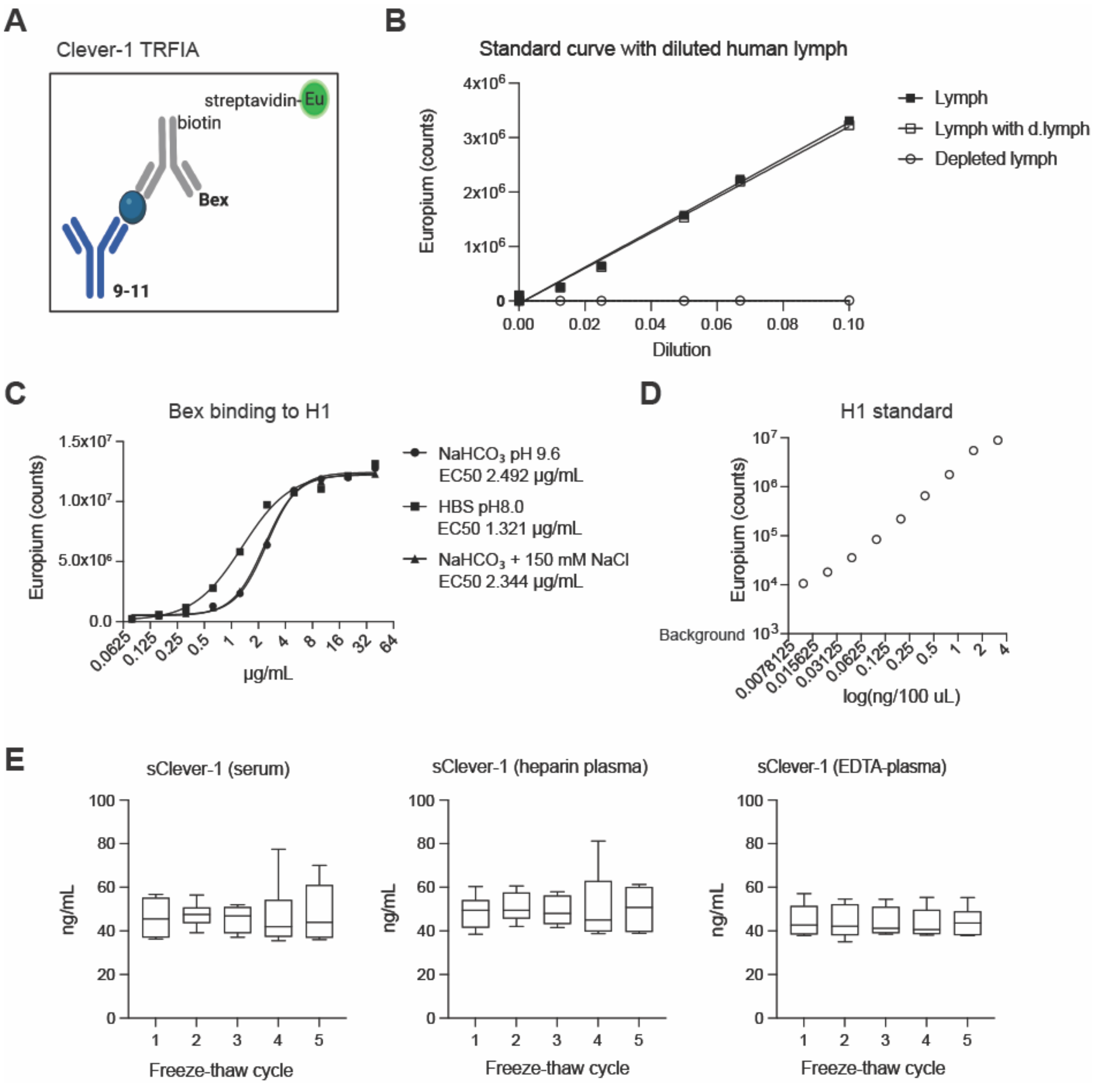

**Supplementary Figure S1. TRFIA assay to measure sClever-1. A**, Schematic of assay design. Blue ball indicates sClever-1 or H1 binding. **B**, Standard curve using serial dilutions of lymph. Clever-1 depleted lymph was used to show the specificity of the signal. **C**, Bexmarilimab (Bex) binding to plate-bound H1. Three different buffers for H1 coating were tested. HBS = HEPES buffered saline. **D**, Standard curve with plasma spiked H1. **E**, Stability of sClever-1 in serum, heparin plasma and EDTA-plasma after 5 freeze/thaw cycles. Related to Figure 1A.

**A**

Correlation between ln-transformed sClever-1 and age

| Group | Healthy control | Breast cancer | MATINS trial |
|---|---|---|---|
| n | 32 | 138 | 193 |
| Pearson's r | 0.14 | 0.22 | 0.0032 |
| *P*-value | 0.45 | ** | 0.96 |

**B**

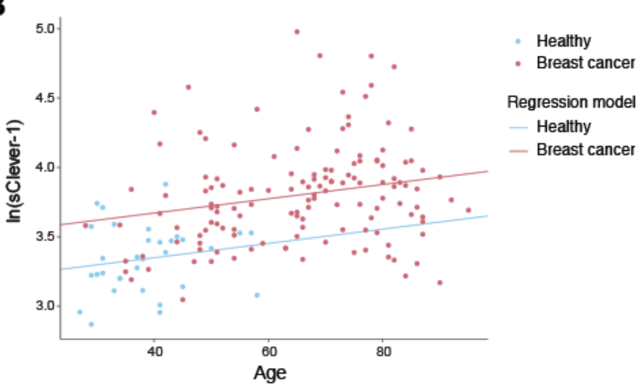

**C**

| Age | Cancer | Predicted sClever-1 (ng/ml) |
|---|---|---|
| 20 | No | 25.6 |
| 20 | Yes | 35.3 |
| 80 | No | 35.0 |
| 80 | Yes | 48.2 |

ln(sClever-1) = 3.14 + 0.0052 x Age + 0.32 x Cancer,
where Cancer = {0, if healthy; 1, if cancer}

**D**

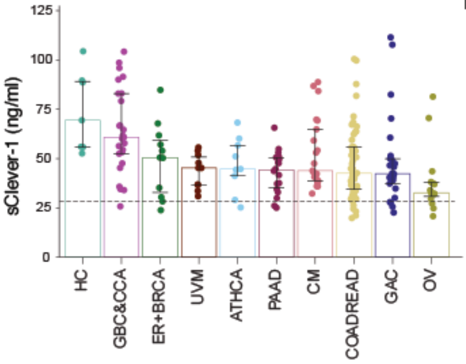

**E**

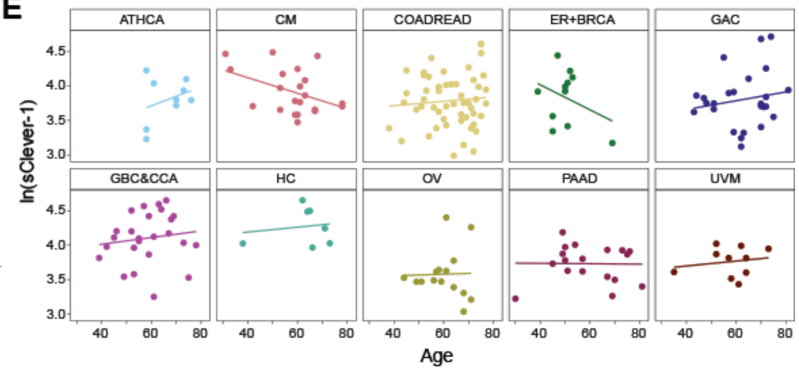

**Supplementary Figure S2. sClever-1 and age. A**, Pearson's correlation coefficients (r) and significance levels for the correlation between ln-transformed sClever-1 and age. **B** and **C**, Multiple linear regression analysis for modeling the effects of cancer and age on plasma sClever-1 levels. A scatter plot showing fitted regression functions for healthy controls and breast cancer patients (**C**), and a table displaying predicted sClever-1 levels in four different cases (**D**). **D**, sClever-1 levels across different cancer cohorts of the MATINS trial (median ± IQR). The dashed line represents median sClever-1 level of healthy controls. **E**, Scatter plots and least-squares-fitted lines showing the association between sClever-1 and age in each cancer cohort of the MATINS trial.

ATHCA, anaplastic thyroid cancer; CM, cutaneous melanoma; COADREAD, colorectal adenocarcinoma; ER+BRCA, Estrogen receptor positive breast cancer; GAC, gastric adenocarcinoma; GBC&CCA, gallbladder cancer and cholangiocarcinoma; HC, hepatocellular carcinoma; OV, ovarian cancer; PAAD, pancreatic ductal adenocarcinoma; UVM, uveal melanoma; sClever-1, secreted Clever; **, $p < 0.01$. Related to Figure 1A.

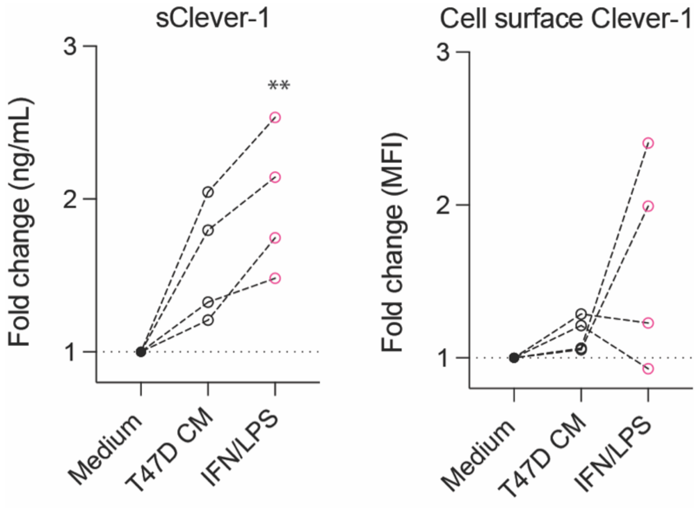

**Supplementary Figure S3. sClever-1 secretion and cell surface Clever-1 levels in macrophages exposed to cancer conditioned medium.** Primary human macrophages (n = 4 healthy donors) were cultured in conditioned medium collected from T47D $ER^+$ breast cancer cell line (ATCC) as previously reported (34) or IFN/LPS for 24h. The sClever-1 was measured from culture medium with the TRFIA assay and cell surface Clever-1 expression (9-11 antibody) by flow cytometry. Friedman test with Dunn's multiple comparison test, ** $P < 0.01$.

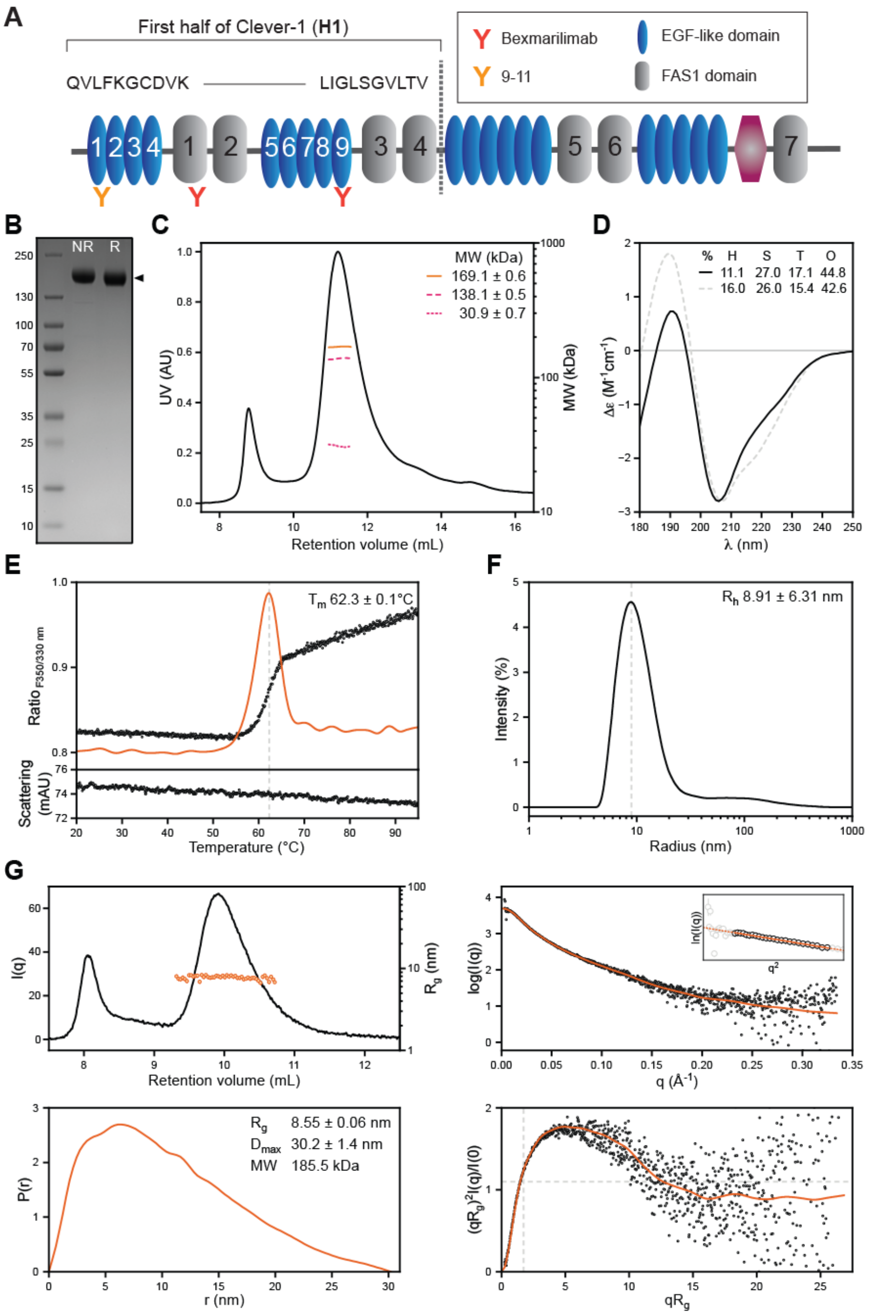

**Supplementary Figure S4. Quality control of recombinant half Clever-1 (H1). A**, The extracellular domain architecture of Clever-1. FAS1, fasciclin-like domain; EGF-like, epidermal growth factor -like; X-link domain in magenta. **B**, Coomassie-stained SDS-PAGE of H1 under reducing (R) or non-reducing (NR) conditions. H1 is denoted by the arrow. Marker sizes are shown on the left in kDa. **C**, SEC-MALS elution profile of H1 (black line) showing the light scattering-derived absolute MW (orange line). The individual protein and glycan components are shown with magenta dash and dot, respectively. **D**, Experimental and simulated far-UV CD spectrum of H1 (black line) and its AlphaFold2 prediction (grey dash), respectively. The average secondary structure contents are denoted by H (α-helix), S (β-sheet), T (turn), and O (other). **E**, Thermal unfolding of H1.

The fluorescence ratio and aggregation signals are indicated in the top and bottom graphs (black circle), respectively. First differential of the fluorescence signal (orange line) indicates the melting temperature ($T_m$). **F**, Distribution of H1 particles sizes (black line). The intensity-weighted average hydrodynamic radius ($R_h$) is indicated by a grey dash. **G**, Baseline-corrected SEC-SAXS elution profile (black line) showing the radius of gyration ($R_g$, orange circles) of H1 derived from a Guinier approximation (top left). Deconvoluted intensity profile (black circle) with a Bayesian fit indicated by an orange line (top right). The inset shows the Guinier fit (orange line) between $0.65 < qR_g < 1.30$ limits (black circle). Pair-distance distribution (orange line) with real-space-derived $R_g$, maximum particle diameter ($D_{max}$), and MW (bottom left). Dimensionless Kratky plot (bottom right) of the intensity data (black circle) and the respective Bayesian fit (orange line). Grey dash reference lines denote the expected peak position of an idealised compact, globular protein.

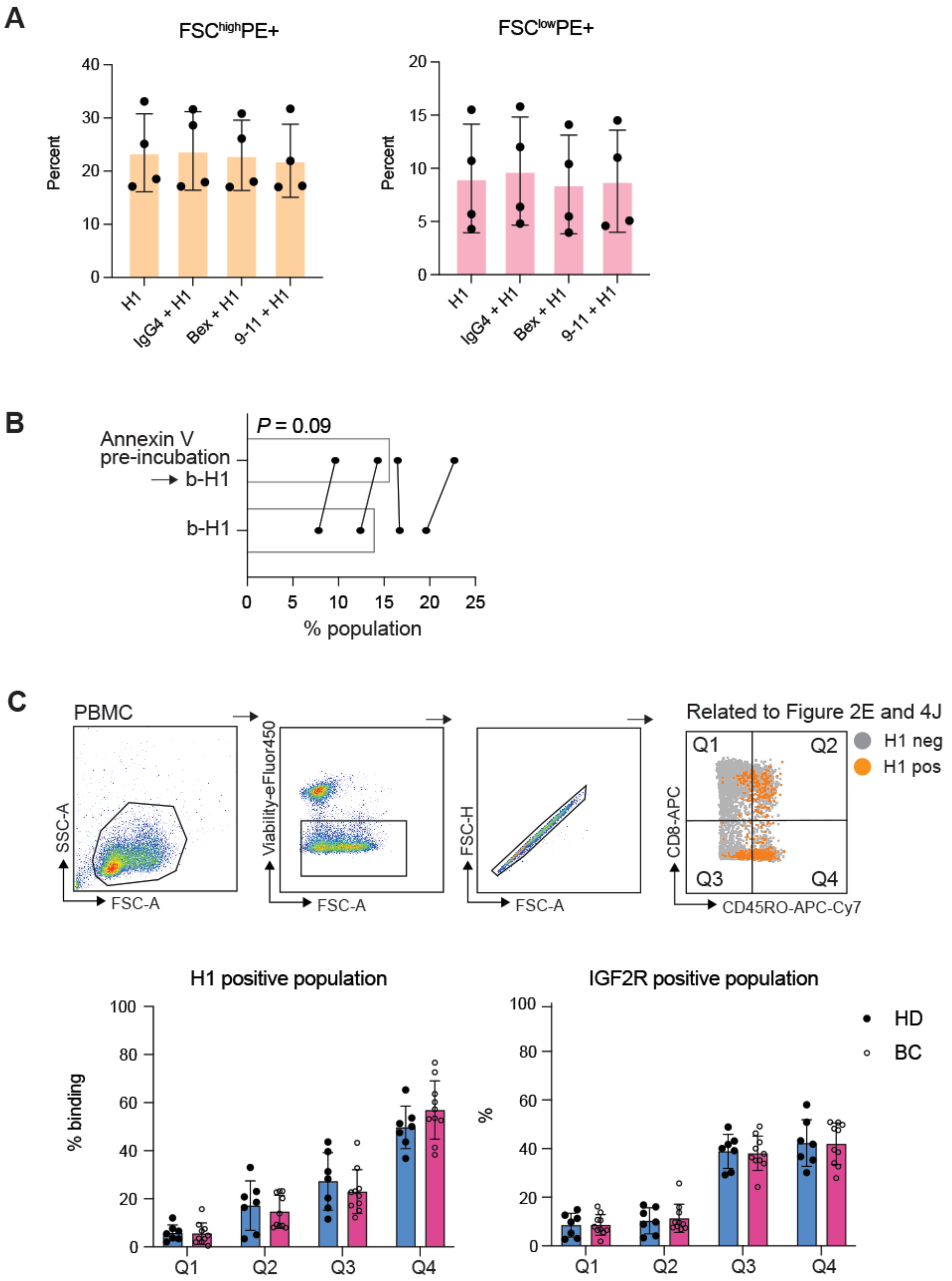

**Supplementary Figure S5. H1 binding on lymphocytes. A,** H1 was pre-incubated with excess amount of anti-Clever-1 antibodies 9-11 and bexmarilimab (Bex) (100 µg/mL) for 30 minutes and H1 binding to T cells was assessed. Related to Figure 2B. **B**, Effect of H1 pre-incubation on Annexin V binding on T cells during ionomycin stimulation. Wilcoxon matched-pairs signed rank test. **C**, Gating strategy for assessment of H1 binding and IGF2R expression on circulating $CD8^+$ memory ($CD45RO^+$) T cells in healthy donors (HD) and breast cancer (BC) patients.

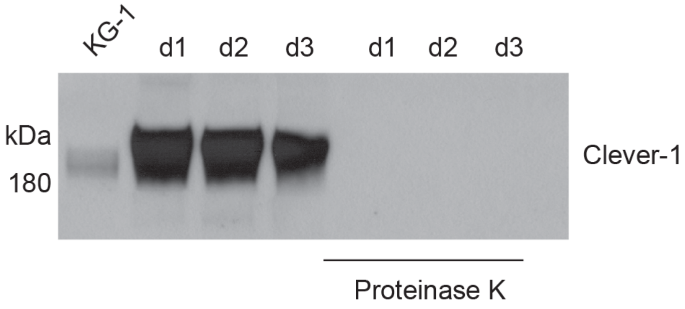

**Supplementary Figure S6. Effect of proteinase K treatment on EV Clever-1 content.** EVs were isolated from plasma of three donors and subsequently treated with or without proteinase K. Clever-1 protein levels were analyzed with Western blot using the 3-372 (InVivo Biotech) mouse anti-human Clever-1 antibody (parent antibody of Bex).

**Supplementary Table S1.**

| Breast cancer patients (n = 138) | | | | |
|---|---|---|---|---|
| Age (mean ± SD) | | 65.5 ± 15.4 | | |
| **Tumor characteristic*** | | **n (%)** | **sClever median (Q25-Q75)** | ***P*-value**** |
| Histolocial subtype | Ductal | 81 (60.9%) | 43.2 (34.9 - 50.4) | |
| | Lobular | 36 (27.1%) | 42.9 (35.1 - 53.5) | 0.94 |
| | Other | 16 (12%) | 40.5 (31.3 - 56.3) | |
| Molecular subtype | ER+PR+ | 100 (75.2%) | 43.7 (36 - 52.5) | |
| | Her2+ | 20 (15%) | 42.1 (31 - 50.9) | 0.33 |
| | TNBC | 13 (9.8%) | 33.7 (31.6 - 50.6) | |
| Grade | I | 6 (4.5%) | 53.2 (47.5 - 56.9) | |
| | II | 90 (67.7%) | 41.9 (34.9 - 50.6) | 0.29 |
| | III | 37 (27.8%) | 43.1 (33.5 - 50.9) | |

* n = 133; five patients with no pathological information available

** Kruskal-Wallis test

**Supplementary Table S2. Multiple linear regression analysis to model the effects of age and cancer on sClever-1 plasma levels.**

| Multiple regression model | | |
|---|---|---|
| n | 138 (breast cancer), 32 (healthy) | |
| F-statistic | 28.42 (DF 2 and 167) | |
| *P*-value | **** | |
| Multiple R-squared | 0.25 | |
| **Explanatory variable** | **Adjusted β (95% CI)[1]** | **P-value** |
| (Intercept) | 3.14 (2.96 - 3.32) | *** |
| Age | 0.0052 (0.0015 - 0.0087) | ** |
| Cancer | 0.322 (0.16 - 0.48) | *** |

[1]Unstandardized β, adjusted for other variables included in the model

β, regression coefficient; CI, confidence interval, **, $P < 0.01$; ***, $P < 0.001$; ****, $P < 0.0001$

### Supplementary Table S3. Characteristics of melanoma and lung carcinoma cohort

| *Characteristic* | *N* | *%* | | | |
|---|---|---|---|---|---|
| **Cancer type** | | | | **Anti-PD-1** | **Anti-PD-L1** |
| Lung | 42 | 72,4 | | 26 | 16 |
| Melanoma | 16 | 27,6 | | 16 | 0 |
| **Lung cancer subtype** | | | | | |
| Adenocarcinoma | 23 | 54,8 | | 15 | 8 |
| Epidermoid carcinoma | 14 | 33,3 | | 8 | 6 |
| Other | 5 | 11,9 | | 3 | 2 |
| **Chemotherapy** | | | | | |
| No | 26 | 44,8 | | 20 | 6 |
| During immunotherapy | 6 | 10,3 | | 6 | 0 |
| After immunotherapy | 26 | 44,8 | | 16 | 10 |
| **Radiotherapy** | | | | | |
| No | 44 | 75,9 | | 32 | 12 |
| During immunotherapy | 5 | 8,6 | | 5 | 0 |
| After immunotherapy | 12 | 20,7 | | 8 | 4 |
| **Lesion removal** | | | | | |
| No | 53 | 91,4 | | 37 | 16 |
| Yes | 5 | 8,6 | | 5 | 0 |
| **Immunotherapy** | | | | | |
| Anti-PD-1 | 42 | 72,4 | | | |
| Anti-PD-L1 | 16 | 27,6 | | | |
| **Gender** | | | | | |
| Male | 36 | 62,1 | | 25 | 11 |
| Female | 22 | 37,9 | | 17 | 5 |
| *Characteristic* | *Median* | *IQR* | | | |
| **Age at immunotherapy** | 66 | 61 - 71 | | 65 (58 - 72) | 69 (64 - 70) |
| **N immunotherapy doses** | 5 | 3 - 9 | | 5,0 (3,0 - 9,8) | 6,5 (3,0 - 8,25) |
| **sClever** | 46,2 | 33,6 - 55,2 | | 46,9 (33,6 - 54,7) | 45,4 (34,9 - 59,5) |
| **B-Leukocytes** | 7,2 | 6,0 - 8,6 | | 7,1 (6,0 - 8,5) | 7,2 (6,1 - 8,6) |
| **B-Basophils** | 0,05 | 0,03 - 0,06 | NA = 6 | | |
| **B-Eosinophils** | 0,18 | 0,09 - 0,28 | NA = 5 | | |
| **B-Lymphocytes** | 1,78 | 1,3 - 2,2 | NA = 6 | | |
| **B-Monocytes** | 0,71 | 0,51 - 0,83 | NA = 6 | | |
| **B-Neutrophils** | 4,1 | 3,1 - 5,3 | NA = 3 | | |
| **CRP (<1 = 0.1)** | 6,5 | 2 - 19,8 | NA = 4 | | |
